# SVPG: A pangenome-based structural variant detection approach and rapid augmentation of pangenome graphs with new samples

**DOI:** 10.1101/2025.07.11.664486

**Authors:** Tao Jiang, Heng Hu, Runtian Gao, Zhongjun Jiang, Murong Zhou, Wentao Gao, Shengming Zhou, Guohua Wang

**Affiliations:** Faculty of Computing, Harbin Institute of Technology, Harbin 150001, China; College of Life Sciences, Northeast Forestry University, Harbin 150000, China

## Abstract

Breakthrough advances in long-read sequencing technologies have opened unprecedented opportunities to study genetic variations through comprehensive pangenome analysis. However, the availability of structural variant (SV) calling tools that can effectively leverage pangenome information is limited. In addition, efficient construction of pangenome graphs becomes increasingly challenging with acquisition of larger number of samples. In this study, we present SVPG, an approach that leverages haplotype-resolved pangenome reference for accurate SV detection and rapid pangenome graph augmentation from long-read sequencing data. Compared to state-of-the-art SV callers, SVPG maintained superior overall performance across different sequencing technologies and coverages. SVPG also achieved notable improvements in calling rare and individual-specific SVs on both simulated and real somatic datasets. Furthermore, in a benchmark involving 20 samples, SVPG accelerated pangenome graph augmentation by nearly 10-fold compared to traditional augmentation strategies. We believe that this novel SVPG method, has the potential to revolutionize SV detection and serve as an effective and essential tool, offering new possibilities for advancing pangenomic research.

## INTRODUCTION

Structural variants (SVs), defined as DNA sequence alterations larger than 50 base pairs (bp), encompass insertions, deletions, inversions, translocations, and duplications^1, 2^. Compared to single nucleotide variants, SVs are fewer in number; however, their broader genomic impact often leads to more significant phenotypic changes^3, 4^. In clinical medicine, SVs are closely associated with numerous genetic diseases, cancers, and complex disorders, emerging as crucial molecular markers for disease diagnosis and treatment^5, 6^. Simultaneously, SVs play an irreplaceable role in revealing genome evolution and analyzing population genetic diversity, providing key insights into species evolution and population characteristics^7^.

SV detection has primarily relied on single reference genome sequences as analytical benchmarks. However, traditional methods such as Sniffles2^8^ and cuteSV^9^ face a fundamental limitation: the single reference genome cannot comprehensively reflect the rich genetic variation within species. This issue is pronounced in highly polymorphic regions or population-specific sequences, often leading to reference bias^10-12^. This bias affects the accurate mapping of sample-specific sequences, thereby reducing SV detection reliability. The emergence of pangenomes offers a new solution to this challenge. By integrating genomic information from multiple representative individuals, pangenomes create a complete genetic variation map encompassing both shared sequences and population-specific variants^13-15^. This comprehensive genetic background significantly improves genome alignment accuracy, offering unique advantages in SV detection, particularly in complex variant regions. Recently, the Human Pangenome Reference Consortium (HPRC) successfully constructed the first human pangenome reference draft through sequencing and assembly of diverse global populations, establishing a solid foundation for improving SV detection accuracy and understanding human genetic diversity^14^.

Beyond improving detection of known variants, *de novo* variant discovery within the pangenome makes capturing individual-specific variants more flexible and pangenome refinement more scalable. By utilizing pangenomes as backgrounds of genetic diversity, researchers can precisely identify rare variants absent from population-level data within a single genome. This fundamentally accelerates the discovery of rare SVs, especially those critical for detecting cancer-specific SVs. Furthermore, as genomic studies expand in scale, pangenome construction and maintenance face substantial computational challenges. When integrating new samples into existing pangenomes, traditional approaches typically require re-execution of expensive *de novo* assembly and complex graph construction, with computational burden scaling superlinearly with sample numbers. In this context, rapidly and accurately discover sample-specific SVs from new samples and dynamically integrate pangenomes without complete reconstruction significantly reduces computational costs and accelerates iterative pangenome updates.

Although the utility of pangenomes for SV detection has been widely recognized, there is a lack of effective tools to fully leverage this advanced resource; existing tools have notable limitations. Early attempts, such as the pangenome-guided SV detection tool PanSVR^16^ and several short-read-based genotypers^12, 17-19^ demonstrate the potential of leveraging pangenomes for accurate SV detection and genotyping. However, these tools were primarily designed for short-read sequencing data, which inherently limits their ability to resolve large SVs. Recent developments pangenome-based tools such as SVarp^20^ and miniSV^21^ represent significant advances in *de novo* SV discovery from pangenome. SVarp can generate SV-related contigs through local assembly, while miniSV provides direct SV calling based on the pangenome graph. Both tools are limited to reporting variants outside the pangenome graph, and their performance in diverse genomic contexts has yet to be evaluated comprehensively.

Here we present SVPG, a tool designed for SV detection and pangenome graph construction. Leveraging the pangenome as a reference background, SVPG accurately filters false-positive calls in germline SV detection and refines SV breakpoint coordinates, improving overall SV detection performance. Additionally, SVPG supports identification of *de novo* SVs from pangenome graphs, capturing low-frequency variants associated with rare disease or cancer-related loci, providing critical support for rare disease etiology research and personalized cancer therapeutics. Finally, novel SVs called by SVPG can be seamlessly integrated back into the pangenome framework, enabling efficient enhancement and continuous iteration of the graph structure, thereby establishing a reciprocal feedback mechanism between variant detection and graph evolution.

## RESULTS

### Overview of SVPG

SVPG integrates two SV detection modes: pangenome-guided and pangenome-based SV detection (Fig. 1). In the pangenome-guided mode, SVPG first converts collected SV signals from a BAM file into signature reads and realigns these reads to the pangenome reference. By thoroughly analyzing the topological features and path transition patterns of SV signature reads in the pangenome graph, SVPG achieves high-precision SV breakpoint localization while optimizing the quality of original feature signals. In the pangenome-based mode, SVPG directly analyzes the alignment characteristics and patterns of raw read sequences within the pangenome graph structure without relying on the linear reference genome, accurately detecting *de novo* SVs within haplotype paths of graph (see Methods for details). This dual-mode design provides SVPG with broader applicability: Researchers can select the first mode conservative and highly accurate germline SV calling, capturing both common and novel variants and supporting whole-genome and population-level genotyping. In contrast, the second mode is suited for detecting sample-specific events in tumor analyses and identifies novel or rare variants that are not encoded by any haplotype in the graph. Beyond the above-mentioned SV detection functions, SVPG integrates a graph augmentation pipeline, allowing researchers to rapidly call pangenome-based SVs from population-scale new samples and implementing graph augmentation functionality in conjunction with pangenome construction tools.

**Fig. 1.**
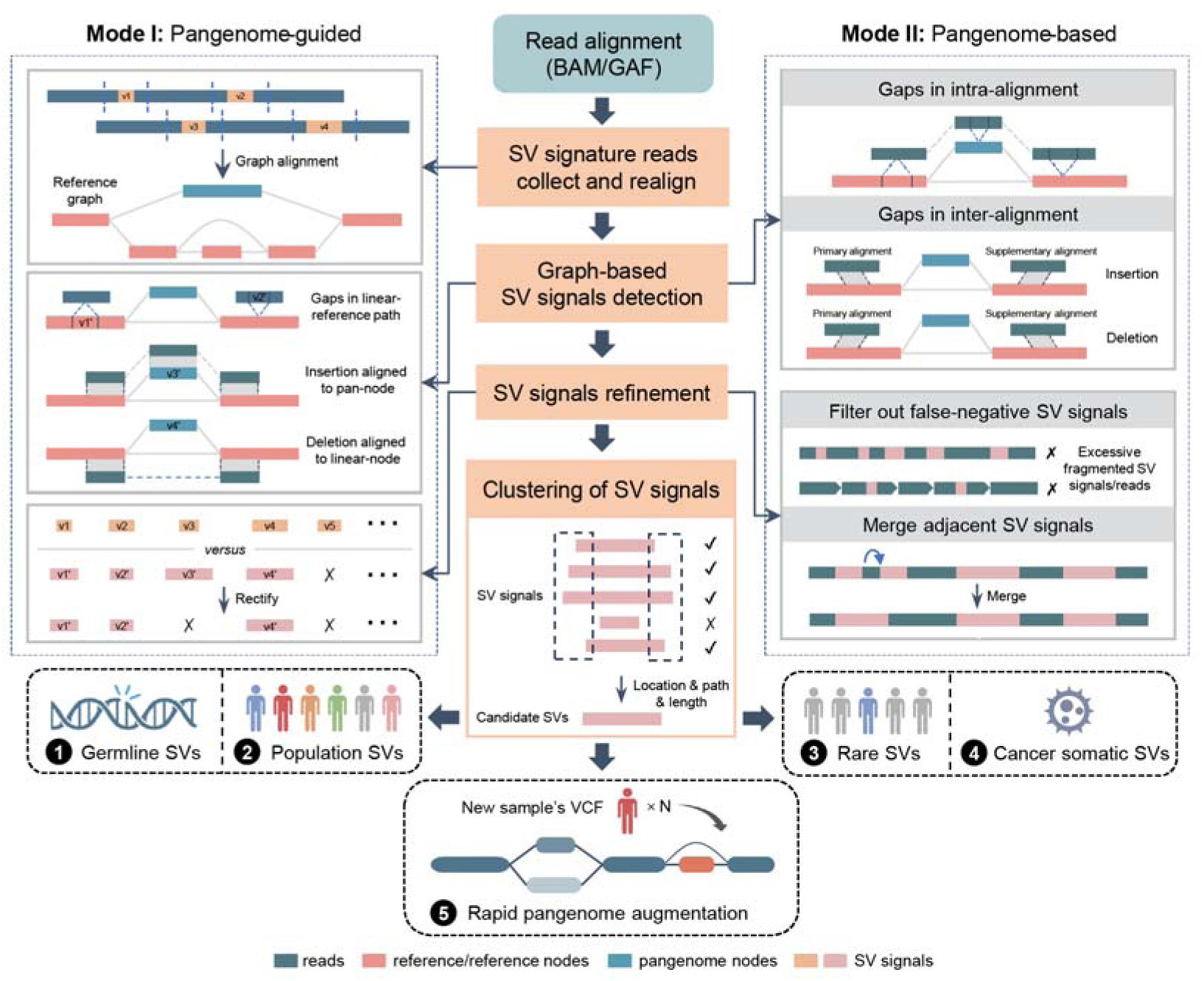
Overview of SVPG. SVPG supports two modes of SV detection, processing either BAM or GAF alignments, which correspond to pangenome-guided and pangenome-based SV detection, respectively. In the BAM-input mode (left), SVPG collects SV signature reads and aligns them to the pangenome reference, followed by graph-aware SV signal extraction and refinement. The GAF-input mode (right) enables direct extraction and refinement of de novo SV signals from sequence-graph alignments. Both modes utilize a common cluster-based refinement step, ultimately generating standard VCF output.

### SVPG enables high performance SV calling on the GIAB ground truth set

We first assessed pangenome-guided mode’s SV calling performance on HG002 sample’s Tier1_ v0.6 ground truth set^22^, from the Genome in a Bottle (GIAB) consortium. We compared the SVPG with five leading long-read linear-reference-based SV callers, including Sniffles2^8^, cuteSV^9^, SVIM^23^, DeBreak^24^ and a recent published from PacBio specific HiFi SV calling tool, Sawfish^25^, and with two pangenome-based callers, miniSV and SVarp. In the high-confidence SV regions, SVPG was among the top performers with F1 scores of 95.8% and 96.6% for ONT and HiFi platforms (see Fig. 2a, Supplementary Table 1 and Supplementary Fig. 1–3 for even more details). In subsequent downsampling datasets, SVPG maintained F1 scores above 90.0% even at low-coverage conditions (ONT-10× and HiFi-5× coverage), consistently demonstrating superior detection robustness.

**Fig. 2.**
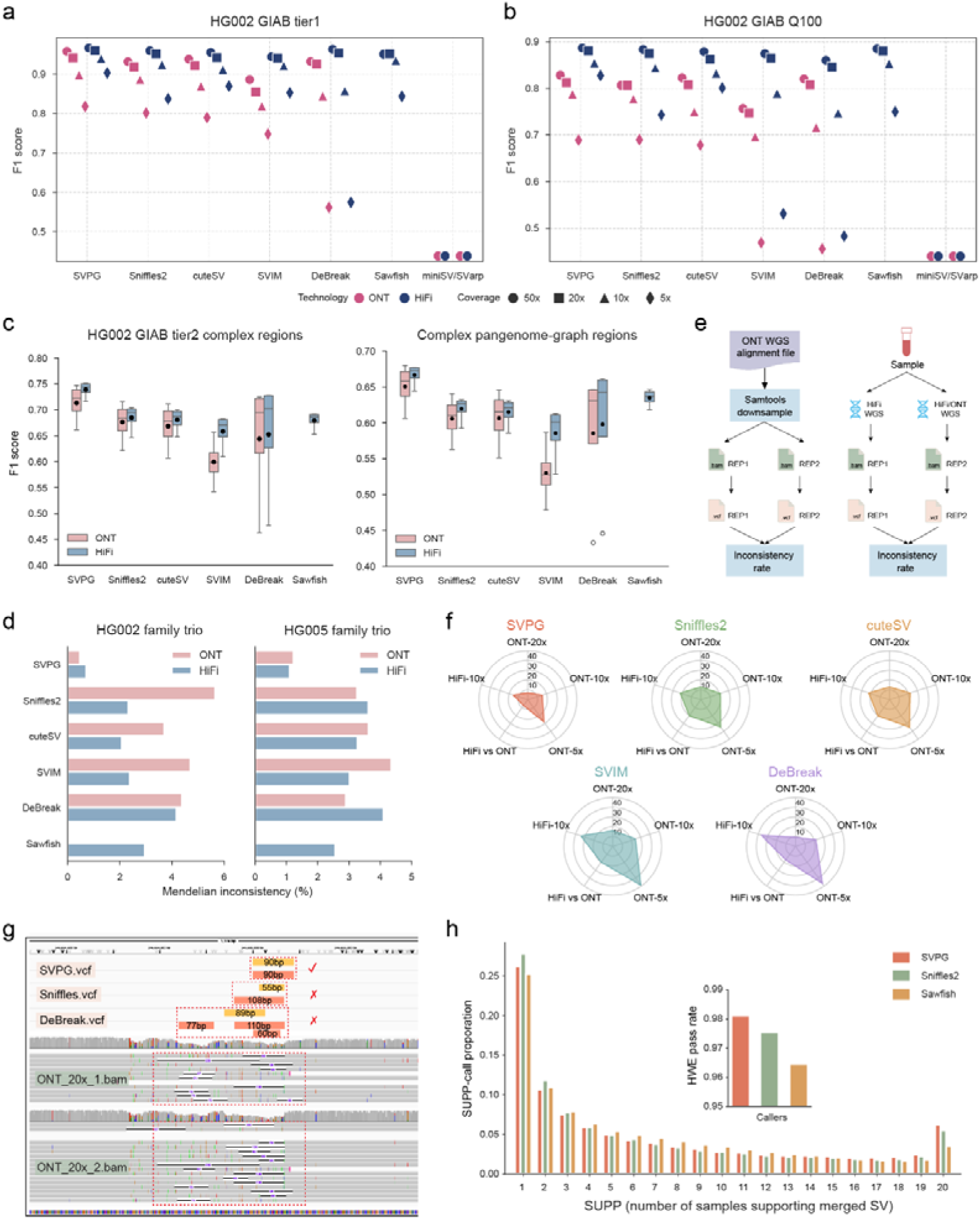
Performance assessment of SVPG in pangenome-guided SV calling. **a** and **b**, Comparison of SV calling F1 score using the GIAB-tier1 and GIAB-Q100 HG002 benchmark across sequencing platform (ONT and HiFi) and coverage. Symbols positioned on the x-axis indicate F1 scores below 0.5. **c**, Comparison of SV calling F1 score at multiple coverages levels (5-50×) across two complex genomic regions on HG002. **d**, Mendelian inconsistency rates across sequencing platform in HG002 and HG005 trio family. **e**, Schematic representation of three replicate-based experiments. **f**, Replicate inconsistency rates for each SV calling tool across five experimental groups: three downsampled ONT replicates from HG002, one cross-platform comparison (HiFi vs. ONT), and one replicate group from HG005 with HiFi 10× coverage. **g**, An example of inconsistent SV calls in the ONT-20× downsampling group. In the top panel, yellow and red bars represent SVs called by SVPG, Sniffles2, and DeBreak in two downsampled replicates, respectively. Only SVPG achieved fully consistent calls. **h**, Multi-sample merging results of SVPG, Sniffles2, and Sawfish across 20 HPRC samples. The x-axis represents the sample support count (SUPP) of the merged SV loci, and the y-axis shows the proportion of SVs at each SUPP level relative to the total number of calls produced by each caller. The inset figure shows comparison of the proportion of SVs that pass the Hardy–Weinberg equilibrium (HWE) test of different tools. The SVs with HWE test scores >= 0.01 were considered pass the test.

We also evaluated all tools using the newly released GIAB HG002-Q100 benchmark set^26^, which provides a next-generation variant reference derived from a fully gapless, telomere-to-telomere genome assembly. On HiFi data, SVPG delivered a leading F1 score of 88.6%, performing on par with Sawfish and outperforming all other methods (Fig. 2b and Supplementary Table 1). In contrast, all tools exhibited a marked drop in performance on ONT data, likely reflecting the substantial number of newly added low-complexity regions in the Q100 benchmark, which pose challenges for technologies with higher base-error rates. Despite this, SVPG achieved the highest F1 score at 82.8%. Analyses of downsampled datasets from both platforms further support above observations. Notably, miniSV and SVarp performed poorly on the HG002 benchmark set primarily because they can only detect SVs outside the pangenome. For this reason, we excluded both tools from subsequent HG002 benchmark evaluations.

Next, we benchmarked performance on representative challenging genome regions: GIAB-released HG002 tier2 regions (containing approximately 6,000 high-confidence SVs in complex or non-consensus sequences), complex graph regions based on the pangenome reference (characterized by high-frequency multi-allelic variants; see Supplementary Fig. 3-4 for distribution, and Methods for construction details), and clinically relevant regions defined in the Challenging Medical Relevant Genes^27^ (CMRG; encompassing SVs in 273 medically significant genes). SVPG demonstrated significant performance advantage over other calling tools when analysis expanded from high-confidence to these complex regions. Particularly, SVPG achieved average F1 scores of 71.4%, 65.1%, and 88.1% across the three challenging regions using ONT dataset with multiple coverages, surpassing the others by 4–12%. For HiFi datasets, SVPG also achieved the highest average F1 scores (Fig. 2c, see Supplementary Table 2-4 for detail). In addition, we applied all callsets to the Q100-defined hard regions to assess the consistency of SVPG’s performance (see Methods for details on region construction). Under this evaluation, SVPG outperformed all other approaches, achieving average F1 scores of 70.2% and 81.0% on ONT and HiFi data, respectively (Supplementary Table 2). Overall, SVPG consistently maintained stable, high-precision performance under various testing conditions, facilitating the reliable discovery of SVs in complex regions.

Given the importance of SV detection in Mendelian disease research, we conducted Mendelian consistency analysis using two representative GIAB-released trio family datasets (Ashkenazim and Chinese Trio). SVPG achieved the lowest inconsistency levels in both families (ONT: 0.5–1.2%, HiFi: 0.7–1.1%; Fig. 2d and Supplementary Table 5). Reduction of SVs showing Mendelian inconsistency represents a crucial step in rapidly identifying *de novo* SVs, as manual screening of all inconsistent variants is typically required when searching for true *de novo* events. We further selected a subset of *de novo* SVs called by SVPG and Sniffles2, in high-confidence regions of HG002 sample for further validation using Integrative Genomics Viewer (IGV)^28^ (Supplementary Note 1). In chromosomes 1–3, results revealed that two out of five *de novo* SVs called by SVPG were confirmed as high-confidence variants. In contrast, of the 13 *de novo* SVs called by Sniffles2, 11 were either false positives or located in noisy regions with split alignments, while the remaining two genuine events were successfully identified by SVPG. These findings confirm that pangenome effectively eliminates Mendelian inconsistencies in SV calling.

### Benchmarking SV call consistency using replicate experiments and cross sample analyses

Traditional SV benchmarking heavily relies on the gold standard datasets released by GIAB, which constrains evaluation comprehensiveness and may introduces systematic biases. The replicate-based evaluation strategy enables the efficient quantification of calling consistency and false positives. The core principle of this strategy is that genuine SVs should be consistently detected across replicates, whereas false positives arising from technical artifacts (e.g., library preparation or sequencing errors) typically appear in single samples^29-31^. Therefore, the more inconsistent events there are in replicate experiments, the higher the proportion of potential false positive SVs in the calls. To avoid sensitivity biases caused by uneven sequencing depth in actual repeat sequencing, we simulated replicate sample pairs with the same coverage by downsampling BAM files of the same sample, using different random seeds (Fig. 2e). We ran SVPG’s pangenome-guided mode and other calling tools on replicate sample pairs of HG002 at 20×, 10×, and 5× coverage from ONT, comparing the inconsistency of the SV sets detected. The results showed that SVPG exhibited the lowest inconsistency in all three downsampling replicate experiments (Fig. 2f, 20×: 7.2%, 10×: 15.5%, 5×: 28.3%; see Supplementary Table 6 for detail; Fig. 2g shows an example), significantly outperforming other tools. As sequencing coverage decreased, the inconsistency rate of all tools increased significantly; this is consistent with the theoretical expectation that insufficient supporting reads at low coverage will lead to an increase in the number of false positives. We then validated the results using a set of 10× coverage raw HiFi sequencing data, similar results were obtained, further confirming the stability advantage of SVPG.

We further assessed the consistency of SV calls across different sequencing platforms using of HG002 HiFi and ONT data samples. Using high-accuracy HiFi data as the control, SVPG showed only an 7.4% discrepancy between platforms, substantially outperforming other tools (e.g., 17.1% of the second-best DeBreak). This exceptional cross-platform consistency of SVPG, despite similar F1 scores among tools on the same platform, highlights the ability to reduce false positives by leveraging pangenome-guided refinement and thereby minimize technology-specific biases.

To validate SVPG’s cross-sample consistency at the population level, we next performed SVPG calling on each of the 20 HPRC samples and merged the results, alongside parallel workflows using Sniffles2 and Sawfish. We used the number of supporting samples per merged site as a proxy for population frequency and quantified the proportion of SVs falling into different support intervals for each method. SVPG exhibited a markedly higher fraction of high-support (high-frequency) variants (Fig. 2h), indicating superior sensitivity to recurrent, reliable SVs shared across individuals. Consistently, Hardy–Weinberg equilibrium (HWE) analysis further revealed that SVPG produced the highest proportion of loci (98.1%; HWE test scores >= 0.01 were considered pass) conforming to expected allele-frequency distributions. This closer agreement with the theoretical HWE model highlights SVPG’s improved genotyping consistency and reduced bias. Overall, these results highlight SVPG’s advantage in population-scale analyses: leveraging the pangenome framework, SVPG enforces unified representations of the same SV across samples, thereby suppressing sample-specific noise and enhancing cross-sample consistency and comparability.

### Pangenome-based rare SV calling performance assessment

Rare SVs predominantly represent recent or *de novo* structural mutations that reveal potential genomic mutation mechanisms. The 1000 Genomes Project demonstrated that 96–99% of variant alleles in the human genome are shared within populations^32^, and this information has now been systematically incorporated into pangenome references. Leveraging this characteristic, we developed a pangenome-based SV calling mode, which established a reliable approach for sample’s rare SV identification by detecting *de novo* SVs that were absent in the pangenome. To assess the approach, we curated 66,197 rare SVs of varying types and sizes from an existing database as the gold standard benchmark set^33^, and generated two 20× coverage simulated datasets (ONT and HiFi) containing these representative rare variants. By using the “*bedtools jaccard*”^34^ command, we compared these SVs with the bubble regions in the pangenome reference, and found that only 2.6% of the regions overlapped, further confirming the rarity of these SVs (Methods for benchmark set construction details).

We input the simulated read alignments against the pangenome reference into SVPG’s pangenome-based calling mode, and compared its performance with that of miniSV (SVarp was excluded because it failed to finish after five days of runtime, while SVPG and miniSV completed within several minutes). SVPG achieved F1 scores of 89.8% and 88.8% on ONT and HiFi data, respectively, surpassing that of miniSV by 8.1% and 4.5%, respectively. SVPG maintained low false-positive rates while successfully recalling more accurate rare SVs (Fig. 3a, Supplementary Fig. 5 and Supplementary Table 7). This performance advantage was consistent across different SV size ranges and SV type, with particularly notable improvements in regions associated with transposable element activity, such as around 300 bp, 2 kb, and 6 kb (Fig. 3b). Additionally, the size distribution of rare SVs exhibited distinct abundance at these intervals, implying that many of these rare SVs may be influenced by the activity of transposable elements (Supplementary Fig. 6). This observation is further supported by our overlap analysis between rare SVs and major classes of repeat elements (Fig. 3c). These results highlight the widespread impact of repeat elements (Alu, LINE1, SVAs) in shaping human genomic diversity, and high-level similarities between our SV catalogue and other studies^1^.

**Fig. 3.**
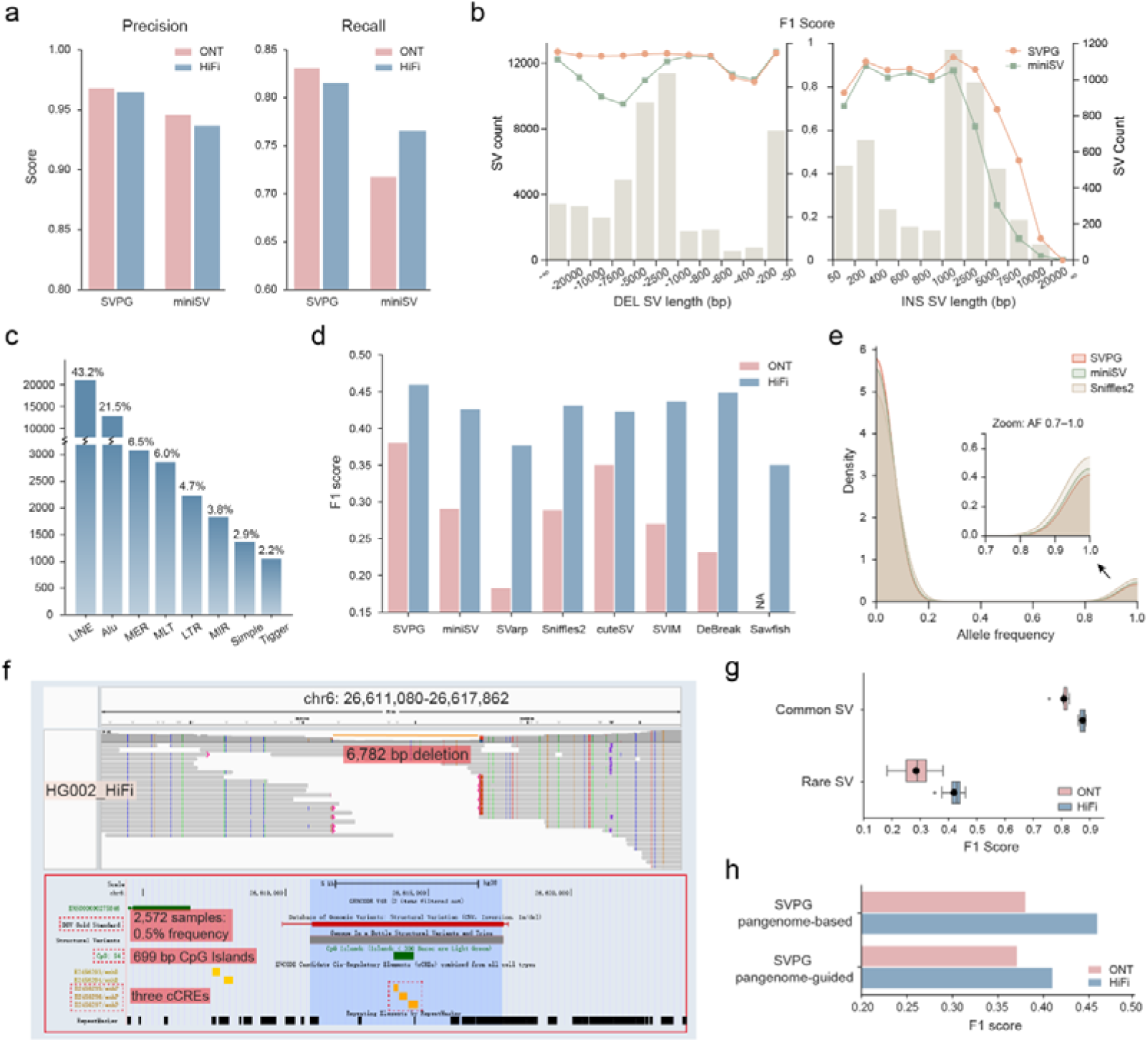
Performance evaluation of SVPG in pangenome-based SV calling. **a**, Comparison of pangenome-based *de novo* SV calling performance on simulated rare SV datasets. **b**, Performance comparison of different rare SV types across different SV length ranges on simulated ONT datasets. The lines indicate F1 score measure, and the bars show the number of SVs called by SVPG. **c**, Overlap analysis between SVPG rare true positive SVs from the ONT dataset and common repeat family. **d**, Performance comparison of pangenome-based tools (top three) and linear-reference-based tools in detecting HG002-Q100 rare SVs. **e**, Allele frequency (AF) distribution of true positive HG002-Q100 rare SVs called by SVPG, miniSV and Sniffles2. The main panel shows kernel density estimates of AF across all variants in the HGSVC dataset. The inset highlights the AF range from 0.7 to 1.0. **f,** A 6,782 bp rare large deletion (chr6: 26,611,080) called by SVPG in HG002. The bottom panel of UCSC Genome Browser snapshot indicates the SV supported by the Database of Genomic Variants (DGV, marked in red), with a population frequency of ∼0.54%, confirming its rarity. The deletion overlaps a 699 bp CpG island and three candidate cis-regulatory elements (cCREs, yellow), suggesting potential disruption to local regulatory activity. **g**, Performance comparison for calling common (six tools) and rare SVs (eight tools) in the HG002-Q100 benchmark set**. h**. Performance comparison of SVPG in pangenome-based versus pangenome-guided modes for HG002 rare SV detection.

We further benchmarked SVPG on HG002 to analyze the performance to detect rare SVs in real sample. Our evaluation was based on two key assumptions: (1) The existing pangenome reference already covered the most common SVs, so *de novo* SVs called from the pangenome graph are likely to represented individual-specific rare SVs; (2) An individual’s rare SVs should be a subset of their complete SV spectrum, with benchmark-validated SVs considered as true positive. Specifically, we stratified the GIAB-Q100 SVs using the pangenome reference: variants not overlapping with pangenome bubbles were labeled as rare SV ground truth, while the remaining were labeled as common SVs. We then aligned HG002 sequencing data to the pangenome reference and performed pangenome-based SV calling (i.e., SVPG, miniSV and SVarp) and linear-reference-based SV calling (i.e., Sniffles2, cuteSV, SVIM, DeBreak, Sawfish). Overall, SVPG achieved 3.0%∼19.8% and 1.0∼10.8% more F1 scores than other callers on ONT and HiFi data from the Q100-rare SVs (Fig. 3d and Supplementary Table 8), respectively. These SVs were further confirmed as genuinely rare through verification against the Human Genome Structural Variation Consortium (HGSVC) dataset of 65 individuals (Fig. 3e). Fig. 3f shows an example of a rare deletion called by SVPG, with the UCSC Genome Browser screenshot suggesting that this variant may be pathogenic.

Moreover, we found that rare SVs are substantially more difficult to detect than common ones (Fig. 3g). Even so, the pangenome-based strategy remains highly effective: compared with the pangenome-guided mode, the full pangenome-based mode enables SVPG to detect rare SVs with markedly higher accuracy (Fig. 3h). It is owing that pangenome-based mode is more sensitive to sequences absent in the pangenome, while pangenome-guided mode inherits biases from the linear reference, limiting its ability in the fully rare SV detection.

### Pangenome-based somatic SV calling performance assessment

To evaluate the potential of pangenomes for cancer-specific SV detection, we applied the SVPG’s pangenome-based mode into a somatic SV discovery pipeline. We expected that aligning reads from both tumor and adjacent normal tissues to the pangenome, rather than to the single linear reference genome, may help reduce false positives caused by reference bias. We assessed SVPG, miniSV, SVarp and five linear-reference-based callers using recently GIAB-released HiFi datasets from the HG008 cancer cell line and its matched control^35^, as well as the widely used COLO829^36^ tumor–normal pair (see Methods for benchmark details). Following the GIAB-recommended benchmarking, SVPG achieved F1 scores of 86.1% on HG008 and 74.3% on COLO829 using Truvari, substantially outperforming other methods (Fig. 4a, Supplementary Table 9). Compared to traditional linear-reference-based callers (such as nanomonsv^37^ and Severus^38^), the pangenome-based SVPG and miniSV, both exhibited significantly lower number of false positive calls. We further evaluated all methods using minda (an independent somatic SV assessment tool developed for Severus), as well as our custom evaluation script specifically designed for SV-type–based assessment of cancer-associated SVs. Fig. 4b shows that SVPG exhibited highly consistent performance across these distinct evaluation tools. In addition, SVPG maintains strong and well-balanced performance across multiple SV types using evaluation scrip (Fig. 4c and Supplementary Table 10). Furthermore, a closer inspection of HG008 dataset the five INDELs missed by SVPG revealed that four were already represented in the pangenome graph, suggesting that they are likely somatic SVs overlapping with common germline SVs (Figs. 4d show one case; see Supplementary Note 2 for all). Overall, we attribute the improvement and low false-positive rate results to SVPG effectively leverages pangenome characteristics to enable precise inference cancer-specific SV.

**Fig. 4.**
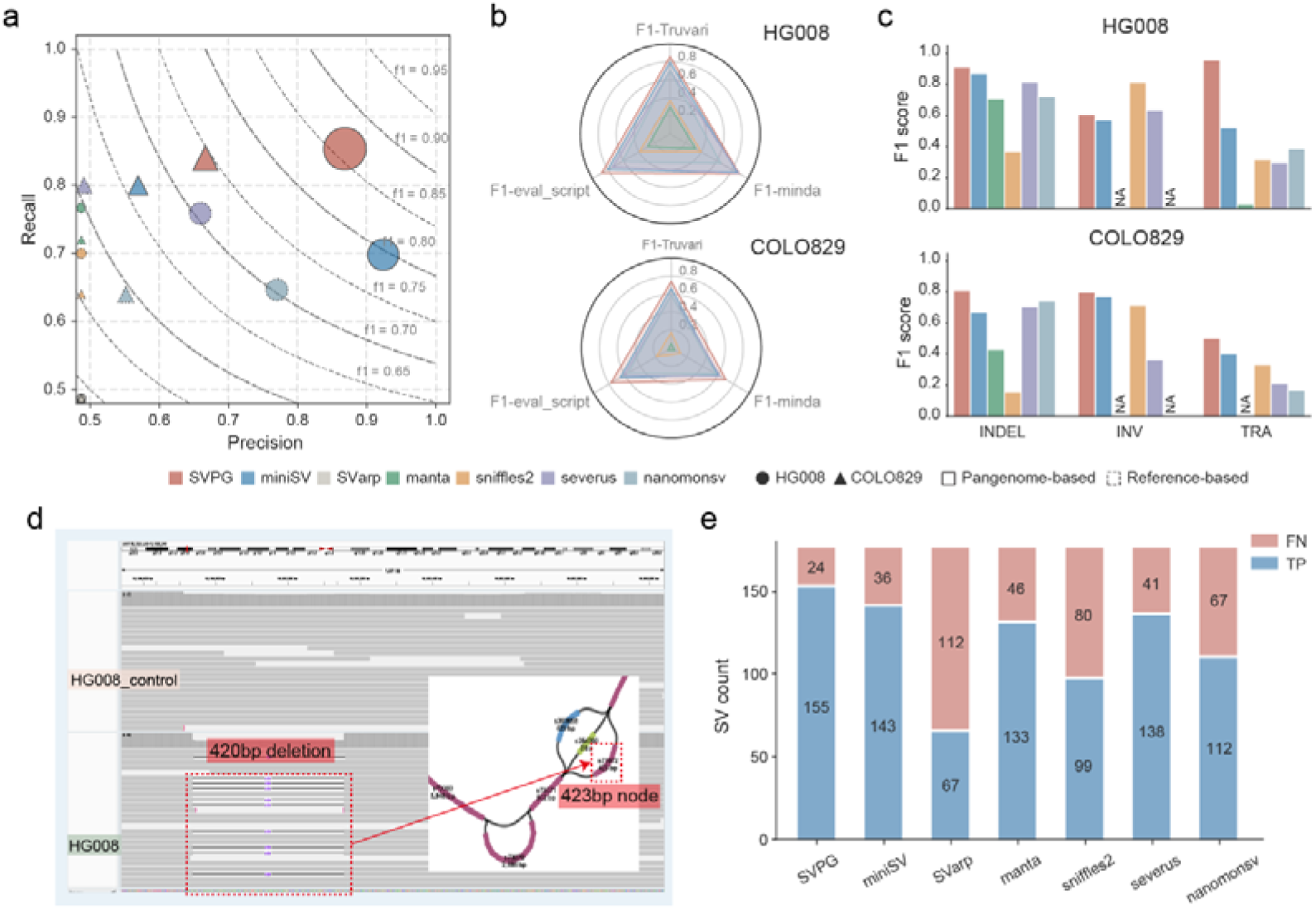
Performance evaluation of SVPG in somatic SV calling. **a**, Precision-Recall bubble plot of somatic SVs called from HG008 and COLO829. Solid lines represent pangenome-based methods and dashed lines represent linear-reference-based methods. Symbols positioned on the axis indicate the score below 0.5. **b**, F1 scores of somatic SV calling on the HG008 and COLO829 assessed using Truvari, minda, and custom evaluation scripts. **c**, F1 score comparison across SV types using custom evaluation scripts. Duplications are treated as insertions and included within the INDEL category. **d,** A missed somatic SV by SVPG in HG008. IGV screenshot shows a 419 bp deletion at chr7: 19,196,736 in HG008, previously reported by GIAB as cancer-specific. Bandage visualization indicates that this deletion is already represented in the pangenome graph, leading to its omission by SVPG. **e**, True positive (TP) and false positive (FP) counts of somatic SV calling on the HCC1395 biologically validated benchmark.

We additionally analyzed an earlier yet widely-studied HCC1395 breast cancer cell line^39^, and its matched normal HCC1395BL. This benchmark was established through the biological validation of a curated subset of SVs derived from consensus germline callsets. Because the dataset lacks rigorously defined high-confidence genomic regions necessary for an accurate assessment of false positives, we prioritized recall as the primary metric for our performance evaluation. As shown Fig. 4e, SVPG achieved the highest sensitivity, identifying 155 out of 179 validated SVs (Supplementary Table 10), followed closely by the pangenome-based method miniSV. We further performed the same evaluation on the consensus benchmark set comprising 1,268 SVs identified by at least two germline SV callers. We found that 951 SVs were not reported by SVPG. Notably, the majority of these SVs were also missed by nanomonsv (949) and Severus (899), indicating that these events may be broadly challenging for current somatic SV callers. Among the 951 SVs not detected by SVPG, cross-validation using *bedtools intersect* showed that 829 overlap with node regions in the GRCh38-90c.r518-based pangenome reference, indicating that these hard-to-call somatic SVs occur in regions enriched for common germline SV (Supplementary Table 11). In contrast, only 25 of the 285 SVs detected by SVPG overlap with pangenome nodes, which is significantly lower than the overlap observed in the unrecalled group (Fisher’s exact test, *p* < 0.001). These regions expose a fundamental limitation of current somatic SV evaluation frameworks, where independent somatic events may overlap common germline variation in highly polymorphic regions. This ambiguity affects SVPG as well as other state-of-the-art callers, underscoring the need for more refined benchmarking strategies that explicitly account for overlapping germline and somatic variation.

### Rapid pangenome graph augmentation from SVPG**’**s graph-called SVs

Traditional pangenome augmentation, exemplified by HPRC efforts, typically relies on complex, time-consuming *de novo* assembly processes which require high-quality sequencing data and multiple refined assembly steps. We propose a rapid graph augmentation strategy based on *de novo* SVs identified by SVPG from graph alignments. By utilizing existing high-quality pangenome reference graphs for initial SV calling and merging, the strategy directly incorporates these variants into the original pangenome, thereby circumventing elaborate assembly procedures. Fig. 5a illustrates the key differences between the two graph augmentation strategies. We selected 20 HPRC samples’ HiFi data and applied SVPG’s graph augmentation mode (Supplementary Table 12), using the HPRC-released original pangenome reference graphs as the foundation for generating augmented versions. Analysis of population-level SVs called by SVPG revealed that these *de novo* SVs, predominantly ranging from 50–500 bp in length, existed at allele frequencies of approximately 0.1, further indicating the rarity (Fig. 5b). Subsequently, we constructed an augmented graph based on *de novo* assembly using the hifiasm^40^ assembler, following the standard pipeline of HPRC. Given our primary focus on SVs exceeding 50bp, all graphs were constructed using minigraph^41^, which is optimized for capturing large-scale genomic rearrangements. Overall, SVPG completed this process within 0.5 day, while for the assembly strategy, even the initial hifiasm assembly step, took up to 3 days, with substantially longer timeframes when including additional refined assembly procedures (Fig. 5c and Supplementary Table 13).

**Fig. 5.**
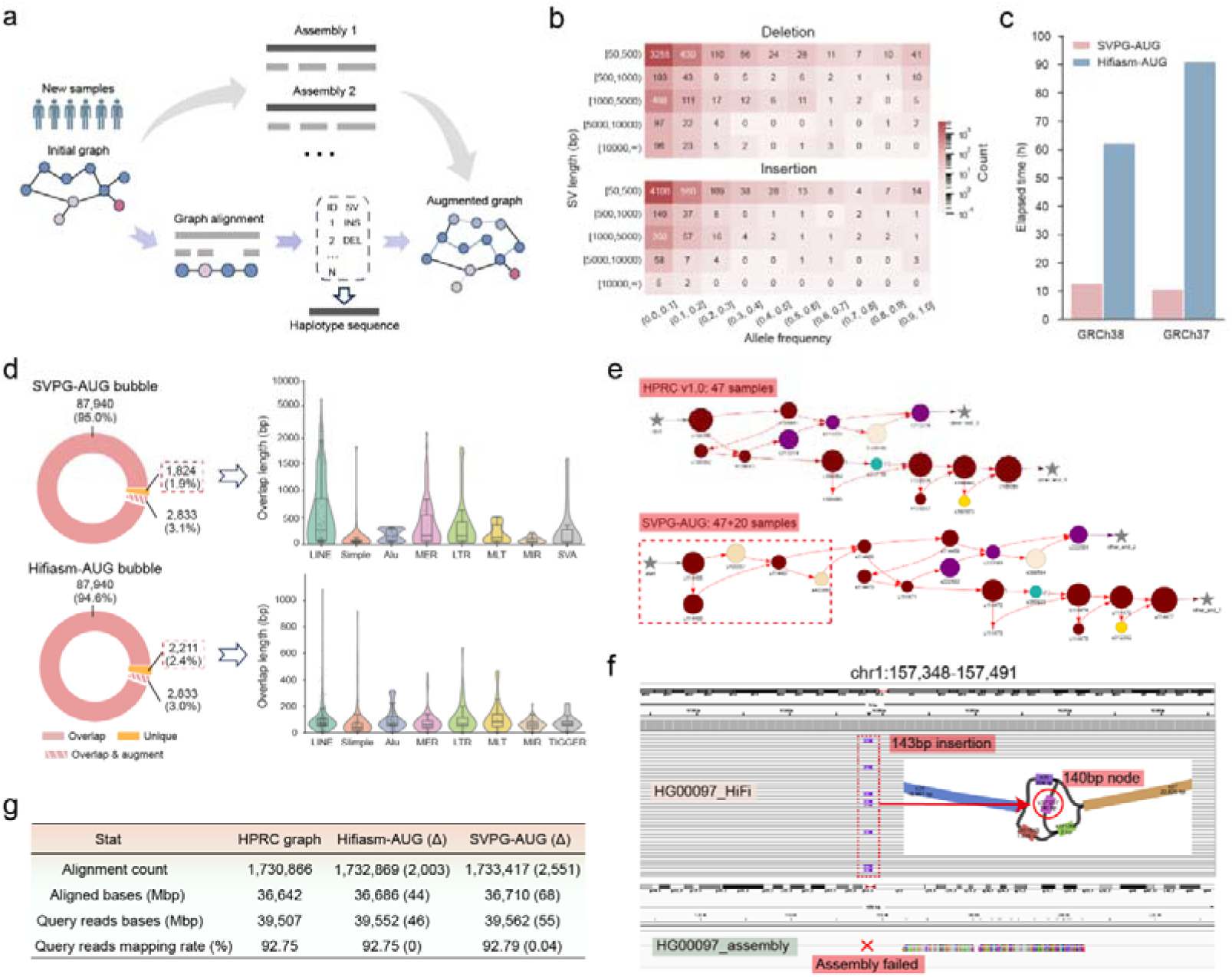
SVPG pangenome graph augmentation results. **a**, Comparison of two approaches to pangenome graph augmentation. The upper grey arrows depict iterative graph augmentation through *de novo* assemblies of new samples (hifiasm-AUG). The lower path shows augmentation through integration of population-scale variant calls from graph alignments (SVPG-AUG). **b,** Distribution of SVs called by SVPG across 20 HPRC samples by SV length and allele frequency. **c**, Comparison of computational time required for constructing SVPG-AUG and hifiasm-AUG. **d**, Composition of bubble regions in GRCh38-based SVPG-AUG and hifiasm-AUG. “Overlap” indicates non-augmented bubbles that are shared between the two augmented graphs (original graph bubbles); “Overlap & augment” represents newly added bubbles compared to the original graph; “Unique” denotes bubbles exclusively found in one augmented graph and not found in the other. The violin plot shows the overlap between unique bubbles and repetitive elements, displaying the top eight repeat families. For SVPG-AUG, overlap lengths >1,000 bp are log-transformed. **e,** Local graph topology at the *MUC6* locus in HPRC GRCh38-based pangenome graph (upper, 17 nodes), and SVPG-AUG (lower, 22 nodes). Additional nodes introduced by SVPG-AUG are highlighted in red box. **f**, An SVPG-AUG-specific node example. The upper IGV panel shows a 143 bp insertion at chr1: 157,348 in HG00097 sample, called by SVPG and successfully integrated into the graph. The lower IGV panel indicates that hifiasm failed to assemble this region. **g,** Summary of alignment metrics obtained by mapping HG002 HiFi reads to the HPRC GRCh38-based pangenome graph and its two corresponding augmented versions. Δ denotes the increase compared to the original graph.

We denote the graph-alignment-based augmented graph as SVPG-AUG and the assembly-based augmented graph as hifiasm-AUG. Compared to the original graph, both augmentation graphs exhibited a substantial increase in nodes and edges (Supplementary Table 14), indicating enhanced the expressive capacity of the graph representations. Analyzing bubble regions which represent variant sites within the graphs, we observed remarkable similarity between the two augmented graphs. Using GRCh38-based augmented graph as an example, we found that 98% of the 92,597 bubbles identified in SVPG-AUG overlapped with bubbles in hifiasm-AUG (Fig. 5d). We further analyzed the subset of bubble regions unique to each augmented graph with respect to annotated repetitive elements. Among the 1,824 SVPG-AUG–specific bubbles, 1,322 (72.5%) overlapped with known repeats, while 1,689 (76.4%) of the 2,211 hifiasm-AUG–specific bubbles overlapped with repeats. Among the eight most abundant repeat families ranked by overlap count (right panel of Fig. 5d), SVPG-AUG exhibited overall greater overlap lengths, particularly with long interspersed elements such as LINEs and Alus. In contrast, hifiasm-AUG displayed a greater number of overlapping regions, especially with shorter, high-copy elements such as SIMPLE repeats. These results indicate that, despite differing methodologies, both augmentation strategies yielded highly consistent variant representations, while also capturing distinct repeat-associated variation.

To compare how pangenomes constructed from complete haplotypes capture specific biologically relevant loci, we focused on the *MUC6* gene region, a locus known for its extensive polymorphism and strong clinical relevance^42^ (see UniProt entry: https://www.uniprot.org/uniprotkb/Q6W4X9/entry). We extracted the subgraph corresponding to *MUC6* regions (chr11: 1,012,823-1,036,718), allowing us to directly inspect how structural diversity is represented across different graph. As shown in Fig. 5e (PanGraphViewer^43^ for graph visualization), SVPG-AUG subdivides an original segment into five distinct nodes, capturing sample-specific structural rearrangements contributed by HG00140 and HG00323 samples. In contrast, hifiasm-AUG maintains a topology similar to the original graph (Supplementary Fig. 9), indicating a more conservative representation of variation in this region. As another case study, we examined 881 genomic regions in the HG00097 sample where hifiasm failed to assemble long-read sequences (all exceeding the 15 kbp average HiFi read length). Within these regions, SVPG identified 25 *de novo* SVs. Fig. 5f presents an example where an SVPG-called SV was successfully incorporated to augment the graph: a 143 bp insertion called by SVPG at the assembly-failed region in the HG00097 sample and encoded in the augmented graph, by decomposing the normal path to create a corresponding bubble structure (s371207). In conclusion, these findings strongly demonstrate that SVPG can effectively augment pangenome structures in assembly-challenging regions, providing more comprehensive genomic characterization for SV analysis.

Finally, we aligned HG002 HiFi data to the pangenome graphs to assess the impact of augmented graphs on downstream applications. Minigraph alignment showed that both augmented graphs improved mapping over the original graph. SVPG-AUG yielded 2,551 more alignment records and 68 Mbp of additional aligned bases, with slightly higher than hifiasm-AUG (Fig. 5g, Supplementary Table 15). Further, we substituted the original graph with augmented graphs to examine their influence on SV calling under the SVPG pangenome-guided mode. Results indicated that the SVPG’s calling performance remained largely consistent across both augmented graphs. In the CMRG regions, SVPG-AUG exhibited modest improvements (Supplementary Table 16). These findings indicate that current pangenome references already capture substantial SV information, and augmented graphs have yet to show marked improvements in common germline SV calling within current benchmarks. However, with advancing technologies and expanding datasets, optimized pangenomes may offer greater benefits, particularly in complex variant regions or diverse population analyses.

## Discussion

This study introduces SVPG, a method to incorporate pangenome information into SV detection of long-read sequencing. Unlike traditional the current long-read SV calling methods that rely on linear reference, SVPG leverages pangenome-guided priors to force-call and annotate widespread common SVs while retaining high sensitivity for sample-specific events. We showed that SVPG consistently outperforms existing state-of-the-art methods, and both unbiased replicate experiments and cross-sample analyses further highlighted its substantially enhanced call consistency in the population scale and markedly reduced false negatives. These findings not only validate the critical importance of pangenome resource in SV detection but also provide a new technical paradigm for future developments in this field.

Furthermore, the SVPG pangenome-based SV calling mode enables *de novo* SV detection from pangenome graph. This offers a novel perspective on rare SV discovery, which traditionally relies on large-scale population sequencing data analysis—an approach that is both resource intensive and prone to missing individual-specific rare variants, particularly without adequate genetic background references. SVPG pangenome-based detection leverages population genetic variation information embedded in pangenomes to achieve accurate rare SV detection at the single-sample level with high sensitivity, eliminating the need for population-scale data support. This capability has significant implications for elucidating disease-associated variants and individual-specific phenotypic mechanisms.

Notably, SVPG demonstrates that pangenome significantly enhances the detection of somatic SVs, the specialized and distinct class of rare SVs. By introducing a pangenome background to effectively filter out the majority of non-cancer-specific somatic SVs, SVPG outperforms linear-reference-based methods with much higher precision. Building on this advantage, SVPG more precisely identified cancer-specific variants through the construction of personalized or population-specific pangenome references from matched normal tissues or cell lines. Further analysis revealed that some false negative SVs unreported by SVPG are actually visible in the germline-derived pangenome, suggesting that current published cancer SV benchmark sets may contain non-genuine cancer-driving mutations. This finding emphasizes that pangenome background is an essential factor when constructing cancer-specific SV benchmark sets.

Our exploration of SVPG application in pangenome graph augmentation also yielded promising results. Using graph SVs called from 20 human samples, we implemented a graph-alignment-based augmentation strategy that integrates SVPG-called graph SVs directly into existing pangenomes. The resulting augmented graphs showed high concordance with those generated from assembly–based approaches, while reducing the computational time required for graph construction by nearly ten-fold across 20 samples, and this efficiency is expected to increase further with larger sample sizes. Notably, while we present several assembly-failure cases, these examples are based on preliminary native assemblies, we acknowledge that with sufficiently accurate and high-coverage reads, together with appropriate post-processing (e.g., trio-binning, polishing, scaffolding), *de novo* assembly can resolve most complex regions, particularly longer segmental duplications or satellite. Nevertheless, when data quality or cost constraints limit assembly success, SVPG provides a cost-effective and scalable alternative for augmenting pangenome graphs, offering substantial practical advantages for large-scale, population-level analyses.

Though this study represents an initial attempt at leveraging pangenomes for SV detection, it still faces several unresolved challenges. First, precise SV detection and reliable assessment in complex genomic regions remain challenging. SVPG performance intrinsically contingent upon the quality of pangenome resources and the accuracy of graph alignment (Supplementary Table 7 shows the impact of different graph-alignment tools on SVPG’s performance). As observed by tools like minigraph, multi-allelic VNTRs pose particular challenges for current graph construction frameworks, preventing an optimal graph representation and alignment. Besides introducing more specific heuristics for complex regions, improving graph-based algorithms remains crucial for advancing pangenome-based SV detection. Second, while alignment-based graph augmentation approaches offer efficient and cost-effective solutions, they rely on an intricate pipeline of multiple tools, including graph alignment, variant calling and merging, and graph construction. This multi-tool dependency introduces compound error sources that are particularly pronounced in multi-allelic regions, where sequence similarities and structural complexities can cascade through each analytical step. In the future, incorporating more fine-grained local assembly may be beneficial for improving sequence accuracy.

As initiatives like HPRC progress and reference graphs continue to evolve, pangenome references are expected to rapidly expand to thousands of haplotypes in the coming years. In this context, the high accuracy and versatility of SVPG position it advantageously for both SV detection and graph augmentation. We anticipate that pangenome-based SV detection methods like SVPG will play an increasingly crucial role in whole-genome analysis pipelines.

## Methods

### SVPG methodology

#### SV signature reads collection and realignment in the pangenome-guided mode

In the pangenome-guided mode, SVPG takes linear reference genome alignment result as the input and extracts SV signatures through two heuristic rules, according to the signal source type: (1) gap sequences in the CIGAR string of primary alignments exceeding predefined thresholds, indicating potential insertions or deletions, and (2) abnormal distances between primary and supplementary alignments from split-read mapping, with sequence lengths exceeding predefined thresholds, usually indicating more complex or larger SVs.

Each detected SV signal is instantiated as an independent object with key attributes: breakpoint coordinates in both reference genome and reads, complete read sequence, read name, and signal source type. SVPG then scans each SV signal object to generate SV signature reads based on their attributes. In the standard pipeline, SVPG extracts an additional 2,000 bp window sequence on both sides of the breakpoints of the SV signal as alignment anchors. For signals from split alignments, the complete original read sequences are used directly. Notably, when SV breakpoints are less than 2,000 bp from read ends, SVPG supplements missing sequences from the reference genome based on the SV signal’s reference breakpoint position, to ensure sequence window completeness. These flanking sequences are concatenated with SV signal sequences to form complete signature read sequences, with key SV signal attributes recorded in the read description line, for subsequent refinement steps.

SVPG employs minigraph for rapid realignment of SV signature reads to the pangenome reference graph. Parameters are configured according to sequencing data types: “*-x asm*” for HiFi data and “*-x lr*” for ONT data. The “*-vc*” parameter enables output of unstable sequence coordinates to support standard GFA graph format.

#### Graph-aware SV signal detection and refinement in the pangenome-guided mode

Unlike BAM files using linear reference coordinates, GAF data structure describes sequence-to-graph alignment information through graph paths and path coordinates. Graph paths comprise ordered nodes and edges, with each node’s start and end uniquely corresponding to linear reference coordinates. SVPG first converts the pangenome graph structure into a dictionary for efficient node querying and graph traversal, with the node name as keys and their properties (sequence length, coordinate information, and topological relationships) as values.

Graph-aware SV signal detection handles two scenarios: (1) Alignment paths containing only linear reference nodes. Here, path discontinuities typically indicate deletions. SVPG accesses the graph dictionary by the missing node names to obtain node coordinates and spans, which accurately represent the SV breakpoints. Additionally, gaps in path CIGAR strings further localize SV breakpoints based on path-start nodes and alignment-start coordinates in the path, thereby capturing variants that are not explicitly encoded in the graph. (2) Alignment paths including non-reference haplotype nodes. SVPG identifies insertions based on haplotype nodes, with variant spans equal to the total span of all haplotype nodes between linear reference nodes. For CIGAR string gaps, processing depends on gap location. Gaps in linear reference node paths are preserved; gaps in haplotype node paths adjust the insertion span represented by that haplotype node. Specifically, insertion-induced gaps increase the original insertion span by the gap length, while deletion-induced gaps decrease it. Finally, for complex paths containing both haplotype and discontinuous nodes, SVPG preserves deletion signals represented by missing nodes, and subtracting haplotype node length from signal span.

The complex topology of pangenome graphs is more prone to alignment artifacts, particularly in highly variable or repetitive regions. To mitigate these effects, SVPG implements stringent signal refinement: all graph-called SV signals within the same signature read are compared against the original SV signature signal for filtering. Specifically, SVPG selects graph-called SVs with breakpoints closest to the raw SV signal. If the graph SV signal span does not meet consistency principles with the raw signal span, all graph SV signals from that signature read are filtered out. The consistency principle determines the reliability of an SV signal based on the span of the called SV (*span__graph_*) and raw SV signal (*span__raw_*); if 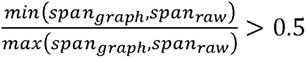, the SV signal is considered highly reliable.

#### Clustering of SV signal

SVPG employs a two-stage clustering strategy to transform scattered graph SV signals into candidate SVs. The first stage performs coarse-grained clustering based on spatial similarity, grouping SV signals of the same type into initial clusters, if their breakpoint distances are within a predefined threshold (default 1,000 bp). This spatial proximity-based clustering broadly captures multiple signals originating from the same SV event.

The second stage focuses on SV signal feature similarity analysis, refining the initial clusters through three feature dimensions: variant span, breakpoint coordinates, and graph path. Span similarity undergoes difference-based normalization, while breakpoint coordinate similarity is normalized using span and parameter σ, where σ is determined by both local cluster SV signal abundance (*localN*) and global SV signal abundance 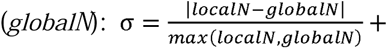 *globalN* Graph path similarity employs Jaccard distance with a penalty µ (default 0.05). The overall similarity between any two SV signals is expressed as: 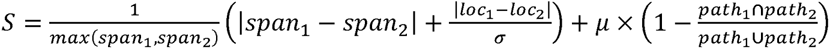. Based on multi-dimensional feature similarity analysis, initial clusters undergo hierarchical clustering (default threshold 0.3) to form highly homogeneous subclusters, each representing potential SV alleles in local genomic regions.

#### SV call and genotyping

Before finalizing candidate SVs, SVPG implements a quality control strategy based on minimum supported reads and alignment path consistency. For each SV cluster, variants sharing identical graph traversal paths are subject to a relaxed read support threshold, effectively recovering high-quality SV signals that may otherwise be missed due to insufficient sequencing depth. This pangenome-based alignment path consistency filtering mechanism effectively balances the sensitivity and specificity of detection.

Clusters passing quality control proceed to final SV calling and genotyping. SVPG uses the median of all SV signal breakpoint coordinates within a cluster as the final candidate SV breakpoint coordinates. Additionally, the SVPG genotyping module assigns optimal genotypes (0/0, 0/1, and 1/1) to each candidate SV through maximum likelihood estimation based on sequence coverage statistics in variant flanking regions.

#### Graph-aware *de novo* SV signal detection and refinement in the pangenome-based mode

SVPG pangenome-based SV detection mode processes GAF format that graphs alignments, converting gap information from each alignment path’s CIGAR string into corresponding SV signals. Besides internal graph alignment information, SVPG encodes SV signals from external graph alignments. Given the lack of explicit supplementary alignment markers in the GAF format compared to BAM format, SVPG sorts multiple alignment records from the same read by genomic position, designating the upstream alignment as primary, and determines SV breakpoint coordinates through primary-supplementary alignment relationships. Subsequently, SVPG infers the SV types (include larger INDELs, inversions, duplications, and translocations) based on the orientation and clipping location of the two segments involved.

Subsequently, SVPG implements post-processing steps to enhance SV signal quality, including: (1) filtering SV signals from reads exceeding thresholds for fragmented SV signals or split operations during alignment; (2) refining SV breakpoints in repetitive regions by identifying locally redundant signals within same genomic windows, and removing inconsistent or spurious SV signals arising from ambiguous alignments in repetitive contexts; (3) merging same-type SV signals’ heterogeneous breakpoints from the same alignments record when their distances fall below a preset threshold (default 500 bp). All SV breakpoints are normalized to a unified coordinate system based on the linear reference, by adjusting for the orientation of their originating graph nodes. Finally, collected SV signals undergo clustering and calling steps identical to the pangenome-guided mode, with results output in VCF format.

#### Graph augment module

SVPG integrates multiple tools to achieve rapid graph augment (Supplementary Fig. 7). Users can provide input through either an organized workspace containing sample sequencing files or a comprehensive file list documenting paths to all sequencing data. For each sample, the workflow begins with minigraph alignment of raw sequencing data to the original pangenome, followed by implementation of the SVPG pangenome-based mode to call *de novo* SVs. Population-level SVs undergo systematic integration through a multi-step process. First, multi-sample VCFs are merged using the “*bcftools merge*”^44^, and redundant variants are eliminated through “*Truvari collapse*”. Subsequently, these SVs are incorporated into the linear reference genome (the initial haplotype used to construct the original pangenome) using “*bcftools consensus*”, generating a haplotype sequence. Before augmentation, uncertain nucleotides (“*N*”) in the sequence are removed to improve alignment efficiency. The resulting haplotype sequence, serving as a carrier for all *de novo* SVs, is ultimately integrated into the original pangenome graph via “*minigraph-cxggs*”, completing the graph augment process.

### Benchmarking methodology

#### Benchmarking SV callers on GIAB ground truths and complex regions

Based on the latest GIAB release, we downloaded HG002 sample raw sequencing reads (HiFi and ONT) alignment data based on the GRCh37 reference genome, and applied SVPG (v1.3), Sniffles2 (v2.3.3), cuteSV (v2.1.1), SVIM (v1.4.1), DeBreak (v1.2), Sawfish (v2.0.3), miniSV (r29) and SVarp (v1.0) to perform SV calling. SVPG, miniSV and SVarp adopts the GRCh37-91c.r559 version pangenome reference release by HPRC as input. SVarp outputs “svtigs” of local assemblies of SV alleles, we aligned these svtigs to the GRCh37 reference using minimap2 and subsequently applied SVIM-asm^45^ to generate a VCF file with the SV breakpoints. Except for Sniffles2, DeBreak and Sawfish, other tools used the same supporting read threshold for low-quality call filtering (Supplementary Note 3 for command lines details).

SV calling results for the HG002 sample were uniformly evaluated using Truvari^46^ (v5.3), against three GIAB-released ground truth sets (HG002_SVs_Tier1_v0.6, HG002_GRCh37_CMRG_SV_v1.00 and GRCh37_HG2-T2TQ100-V1.1), with the “--*includebed*” parameter specifying high-confidence regions and complex genomic regions. Complex graph regions were created using custom scripts. The script traverses the pangenome reference graph along linear reference genome coordinates with a 5,000 bp window size, counting the number of nodes in each window. If the node count >3, the window-covered region is defined as a complex graph region. Q100-hard regions were constructed by subtracting the SV v0.6 tier 1 regions from the Q100 confident regions using the *“bedtools subtract”* command. The remaining genomic intervals, which are not covered by v0.6 tier 1 but still within the Q100 confident regions, were designated as harder evaluation regions.

#### Mendelian inconsistency analysis

We selected two family datasets, Ashkenazim and Chinese family trio datasets for Mendelian inconsistency analysis. Ashkenazim trio sample data directly used GIAB-released HiFi alignment data. Chinese trio samples lacked GIAB-provided HiFi platform alignment results; therefore, we used minimap2 (parameters: “*-ax map-hifi*”) and pbmm2 (Sawfish strictly requires pbmm2 result as input) to align raw HiFi reads to the GRCh37 reference genome. In the evaluation, we selected all SV records >50 bp from the VCF file of each family for consistency verification. Specifically, for the same SV record within the family (with breakpoint distance <1,000 bp), we extracted genotype trios (offspring, father, and mother) and verified their consistency according to Mendelian inheritance rules. For example, when the genotypes of both parents are 0/0, the genotype of the offspring must be 0/0; when one parent is 0/0 and the other is 1/1, the genotype of the offspring must be 0/1. Finally, we calculated the Mendelian inconsistency rate by dividing the number of inconsistent records by the total number of records.

#### Benchmarking the performance of SV caller using replicates

We assessed the inconsistency of replicate experiments using multiple datasets from HG002 and HG005 samples. For the 48× ONT alignment data of HG002, we performed downsampling using “*samtools view*” with default and 99 random seeds to construct paired datasets at 20×, 10×, and 5× coverage. For the HG005 sample, we selected two independent 10× coverage HiFi sequencing datasets and aligned the raw reads to the GRCh37 reference genome using minimap2. In the reproducibility assessment, we employed scripts similar to those used in Mendelian consistency analysis, extracting genotypes from control and case groups for each identical SV record and checking their concordance. It is important to note that inconsistency does not always imply incorrect SV calls. Different callers may report the same true SV with slightly shifted breakpoints or alternative representations, especially in repetitive or microhomology regions. To reduce the risk of misclassifying such cases as false positives, we allowed a breakpoint tolerance of 500 bp and required a size concordance ratio (min/max) of at least 0.7 when comparing SVs across replicates. Only when both conditions were violated did we classify events as inconsistent. Finally, the inconsistency rate was calculated by dividing the number of discordant records by the total number of records. Additionally, we incorporated SURVIVOR^47^, a third-party SV merging tool, to calculate Jaccard similarity for further validation of our assessment accuracy. Specifically, we merged VCF files from two replicate experiments using parameters “*1000 2 1 1 0 50*” and “*1000 1 1 1 0 50*”, then computed Jaccard similarity (A/B) based on the intersection (A) and union (B) of SV records from both files.

#### Benchmarking SV call consistency across 20 samples

HPRC has released raw sequencing data for over 300 samples. From this dataset, we selected 20 samples that were not included in the draft pangenome reference. The HiFi raw reads were aligned to the GRCh38 reference genome using minimap2 and then processed with SVPG in pangenome-guided mode and Sniffles for SV calling; for Sawfish, we utilized pbmm2 alignments to meet the tool’s specific requirements. SV call results across the 20 samples were subsequently merged using Jasmine^48^. For each merged site, the number of supporting samples was extracted from the “*SUPP”* field of the merged VCF, and the proportion of calls for each “*SUPP”* value (*SUPP_call_count / total_call_count*) was calculated. As for the proportion of HWE pass-rate, it was calculated using “*bcftools fill-tags*”.

#### Rare SV detection benchmarking in simulated data

To assess rare SV detection performance, we designed a pipeline to extract 66,197 rare SVs^33^ from the National Human Genome Research Institute (NHGRI) repository to simulate a “super-variant”. Initially, we filtered 95,000 rare indels (duplications were treated as insertions) from the original SV callset using two criteria: “*NSAMP*” and “*NFAM*” = 1, identifying SVs unique to single individuals or families among 14,623 individuals. Subsequently, we removed overlapping SVs, randomly added base sequences of corresponding variant lengths to <ALT> fields of insertion, and for deletions, retrieved base sequences from reference genome breakpoint positions for the fields. Following this preprocessing, we established 66,197 independent rare SVs as our validation benchmark. We then integrated these SVs into the GRCh38 reference genome using “*bcftools consensu*s” to construct the “super-variant”. Based on this variant reference sequence, we simulated 20× coverage ONT and HiFi raw sequencing reads using PBSIM3^49^, aligned them to the pangenome reference (GRCh38-90c.r518 version) using minigraph, generating graph alignment results. Finally, these alignments were input to SVPG, miniSV and SVarp for SV calling; evaluation result from the Truvari summary output files.

#### Somatic SV detection benchmarking

We obtained high-coverage HiFi sequencing data for HG008 and its matched control from GIAB and aligned them to GRCh38-90c.r518 version pangenome reference using minigraph. The resulting graph alignment files underwent independent SV calling through SVPG. Subsequently, we identified cancer-specific SVs through subtracted the normal sample SV calls from the tumor sample calls via our custom script, which filtered VCF files according to defined overlap criteria: SVs in the cancer sample were classified as cancer-specific if their breakpoint did not overlap with that of a normal sample SV within a 1,000 bp window; otherwise, it was considered non-specific. For miniSV, we followed the “tumor-normal pairs” mode as documented on GitHub to identify specific SVs. For SVarp and Sniffles2, we employed the same SV subtraction procedure as SVPG. Manta^50^, severus and nanomonsv directly generated tumor-specific SVs from paired tumor-normal alignment files. Overall performance assessment employed Truvari and minda (commit 182fbcf), while SV-type–based evaluation was performed using a custom evaluation script. The script categorizes variants according to the SVTYPE field in the VCF and currently evaluates three primary SV classes: INDELs (with duplications mapped to insertions), inversions, and translocations. Matching criteria and thresholds are defined to be consistent with those used by Truvari. Supplementary Table 10 shows the custom script show high concordance with Truvari-based evaluations. Before assessment, VCFs with double-entry BNDs were preprocessed to convert them to single representation. For the COLO829 and HCC1395 samples, the calling workflow was performed in the same pipeline as for HG008.

#### Graph augmentation benchmarking in 20 HPRC samples

We used the HiFi raw reads from the 20 HPRC samples mentioned above and input them into the SVPG graph augment mode. For *de novo* assembly-based graph augment, an automated pipeline was employed: hifiasm was used to assemble haplotype-resolved sequences for the 20 samples, which were then iteratively integrated into the original pangenome using minigraph. Basic statistics for graphs were compiled using “*gfatools stat*”, with bubble overlap counts determined using “*bedtools intersect*” (with “*gfatools bubble*” converting graph bubbles into BED format). Additionally, HG002 HiFi sequencing data was aligned to both original and augmented graphs, and alignment statistics were generated using custom script.

## Data availability

### Sequencing and alignment data

GIAB HG002 HiFi data publicly available at https://ftp-trace.ncbi.nlm.nih.gov/giab/ftp/data/AshkenazimTrio/HG002_NA24385_son/PacBio_HiFi-Revio_20231031/. HG002 ONT: https://ftp-trace.ncbi.nlm.nih.gov/giab/ftp/data/AshkenazimTrio/HG002_NA24385_son/UCSC_Ultralong_OxfordNanopore_Promethion/. Ashkenazim trio: https://ftp-trace.ncbi.nlm.nih.gov/giab/ftp/data/AshkenazimTrio/. Chinese trio: https://ftp-trace.ncbi.nlm.nih.gov/giab/ftp/data/ChineseTrio/. HG008 HiFi: https://ftp-trace.ncbi.nlm.nih.gov/giab/ftp/data_somatic/HG008/Liss_lab/PacBio_Revio_20240125/. HG008 Illumina: https://ftp-trace.ncbi.nlm.nih.gov/giab/ftp/data_somatic/HG008/Liss_lab/BCM_Illumina-WGS_20240313/. HCC1395 and COLO829: https://www.pacb.com/connect/datasets/. 20 HPRC samples: https://s3-us-west-2.amazonaws.com/human-pangenomics/index.html?prefix=working/HPRC/.

### Benchmark sets

HG002 genome wide: https://ftp-trace.ncbi.nlm.nih.gov/ReferenceSamples/giab/release/AshkenazimTrio/HG002_NA24385_son/NIST_SV_v0.6/ and https://ftp-trace.ncbi.nlm.nih.gov/ReferenceSamples/giab/data/AshkenazimTrio/analysis/NIST_HG002_DraftBenchmark_defrabbV0.019-20241113/. HG002 medical regions: https://ftp-trace.ncbi.nlm.nih.gov/ReferenceSamples/giab/release/AshkenazimTrio/HG002_NA24385_son/CMRG_v1.00/. Rare SV: https://github.com/hall-lab/sv_paper_042020/blob/master/Supplementary_File_1.zip. Repeats r egion: https://genome.ucsc.edu/cgi-bin/hgTables. HGSVC3: https://ftp.1000genomes.ebi.ac.uk/vol1/ftp/data_collections/HGSVC3/release/Variant_Calls/1.0/GRCh38/. HG008: https://ftp-trace.ncbi.nlm.nih.gov/ReferenceSamples/giab/data_somatic/HG008/Liss_lab/analysis/NIST_HG008-T_somatic-stvar_DraftBenchmark_V0.3-20250220/. COLO829: https://doi.org/10.5281/zenodo.13917379^51^. Consensus benchmark set of HCC_1395_: https://static-content.springer.com/esm/art%3A10.1186%2Fs13059-022-02816-6/MediaObjects/13059_2022_2816_MOESM4_ESM.xlsx. Validation benchmark set of HCC_1395_: https://static-content.springer.com/esm/art%3A10.1186%2Fs13059-022-02816-6/MediaObjects/13059_2022_2816_MOESM5_ESM.xlsx.

### Reference genome

GRCh38: http://ftp.1000genomes.ebi.ac.uk/vol1/ftp/technical/reference/GRCh38_reference_genome/. GRCh37: http://ftp-trace.ncbi.nih.gov/1000genomes/ftp/technical/reference/phase2_reference_assembly_sequence/hs37d5.fa.gz. GRCh38-90c.r518 and GRCh37-91c.r559: https://zenodo.org/records/10693675^52^.

The VCF outputs of all tools, augment pangenome graph, and benchmark scripts are a vailable at https://zenodo.org/records/18456502^53^.

## Code availability

Source code for SVPG is available at available at https://github.com/coopsor/SVPG.

## Author contributions

G.W. conceived and managed the work. T.J. and H.H. designed the method, and H.H. implemented it. R.G., and Z.J. contributed to the experimental evaluations. M.Z., W.G., and S.Z. contributed to data visualization and interpretation. H.H. and T.J. drafted the manuscript. All authors reviewed, revised, and approved the final version of the manuscript.

## Competing interests

The authors declare no competing interests.

## Acknowledgements

This work has been partially supported by the National Natural Science Foundation of China (62225109, 62472120, 62450112). We would like to acknowledge Associate Researcher Yadong Liu, Postdoctoral Fellow Zhendong Zhang and Dr. Shuqi Cao for their valuable suggestions.

## Notes

### Competing Interest Statement

The authors have declared no competing interest.

### Summary of Updates

Modified the author list to be consistent with the manuscript

