## Supplementary File for "SVPG: A pangenome-based structural variant detection approach and rapid augmentation of pangenome graphs with new samples"

**Supplementary Figures**

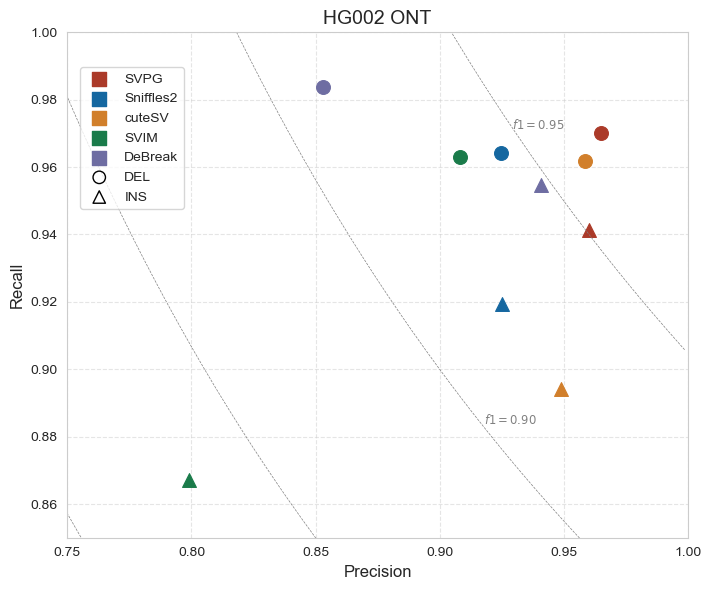

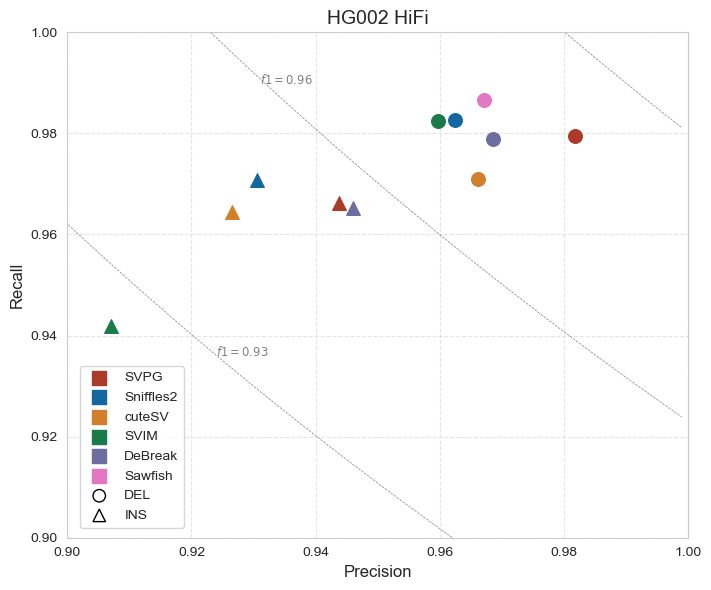

**Supplementary Fig 1.** SV calling performance across different SV types on HG002 47× ONT and 48× HiFi datasets. Measured by using the Truvari for the evaluation of each caller on the GIAB-tier1 ground truth. Sawfish only supports HiFi data, while miniSV and SVarp were excluded due to their poor performance in the GIAB benchmarking.

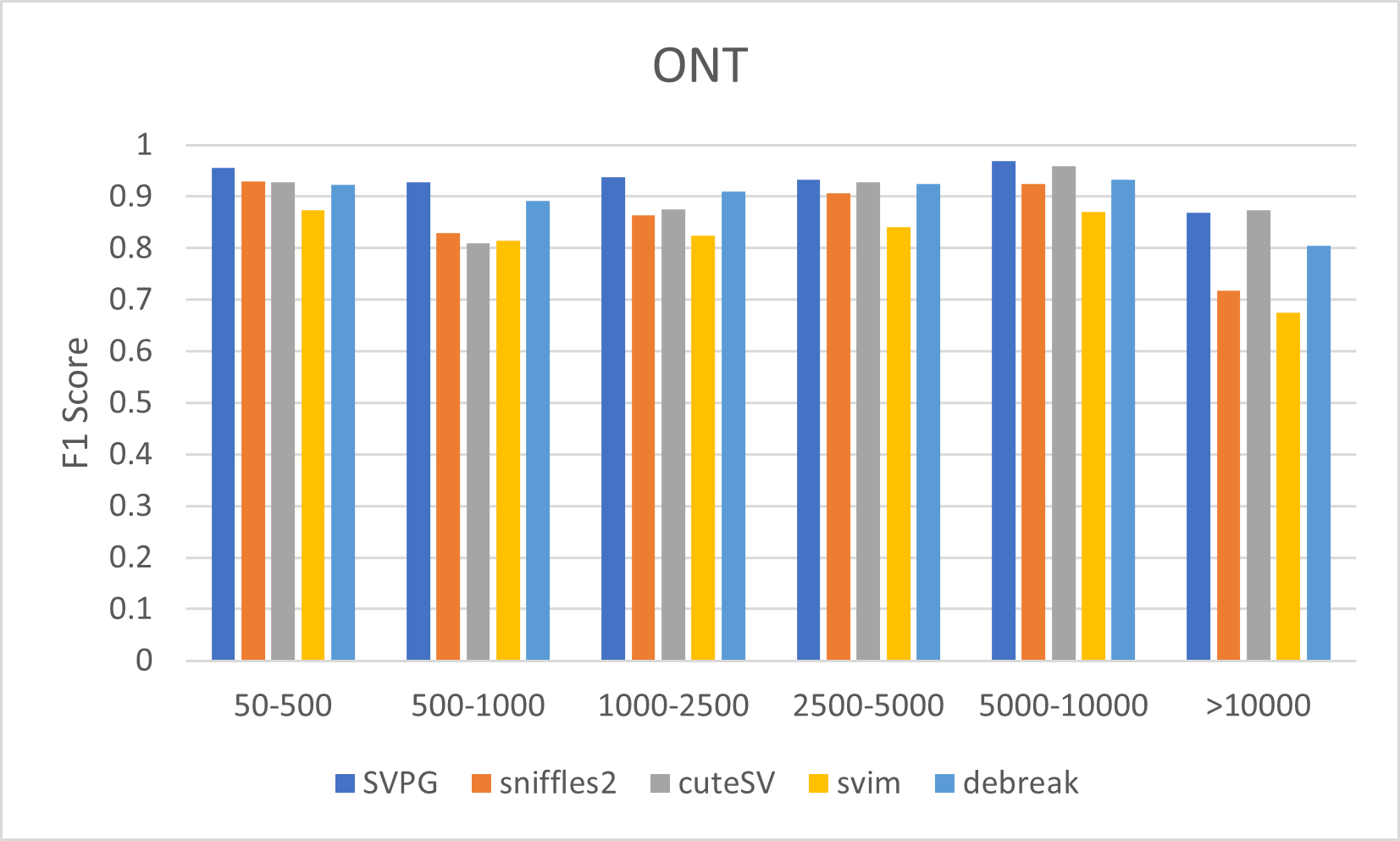

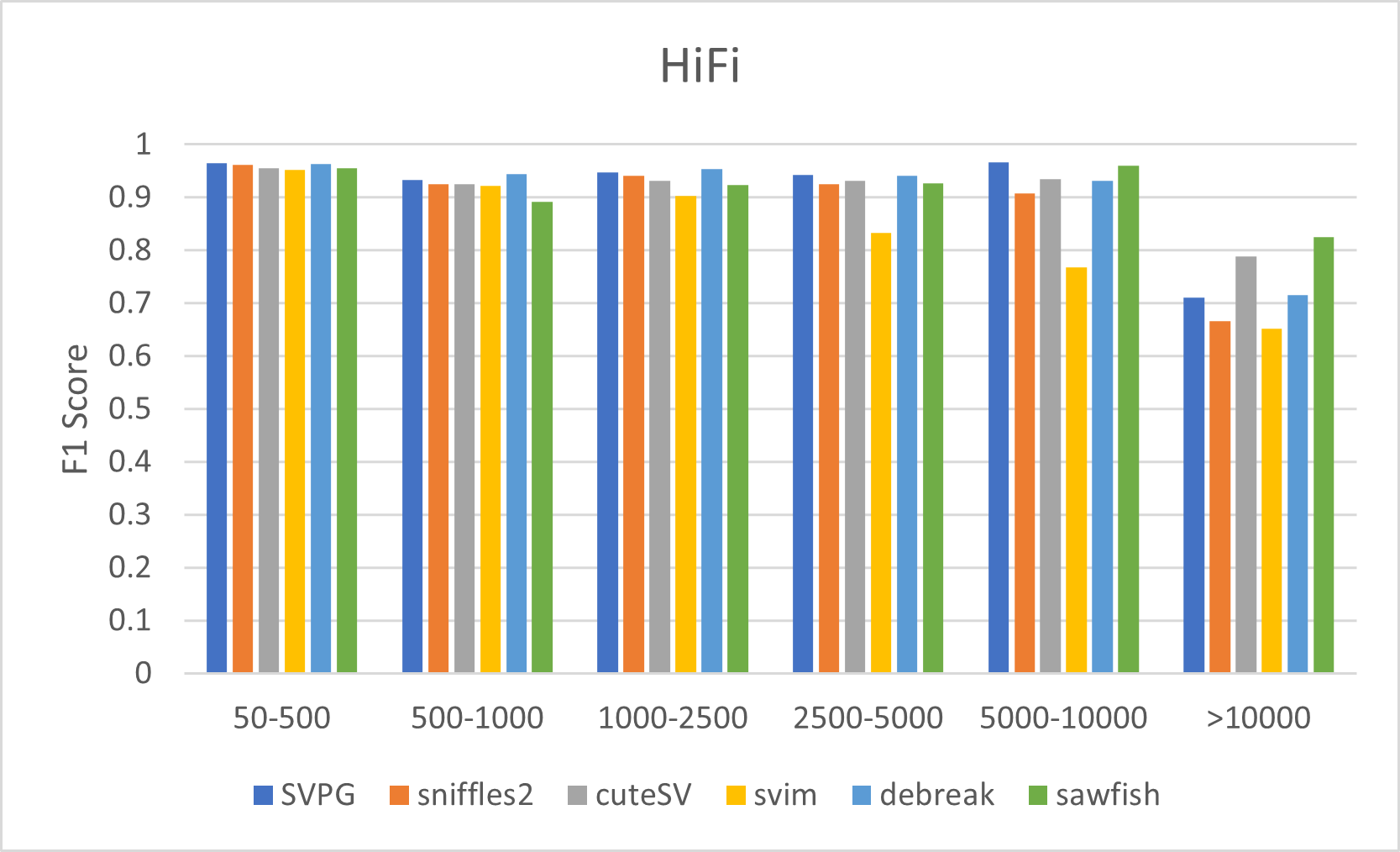

**Supplementary Fig 2.** F1 measure across different SV length on HG002 47× ONT and 48× HiFi datasets. Measured by using the Truvari for the evaluation of each caller on the GIAB-tier1 ground truth. Sawfish only supports HiFi data, while miniSV and SVarp were excluded due to their poor performance in the GIAB benchmarking

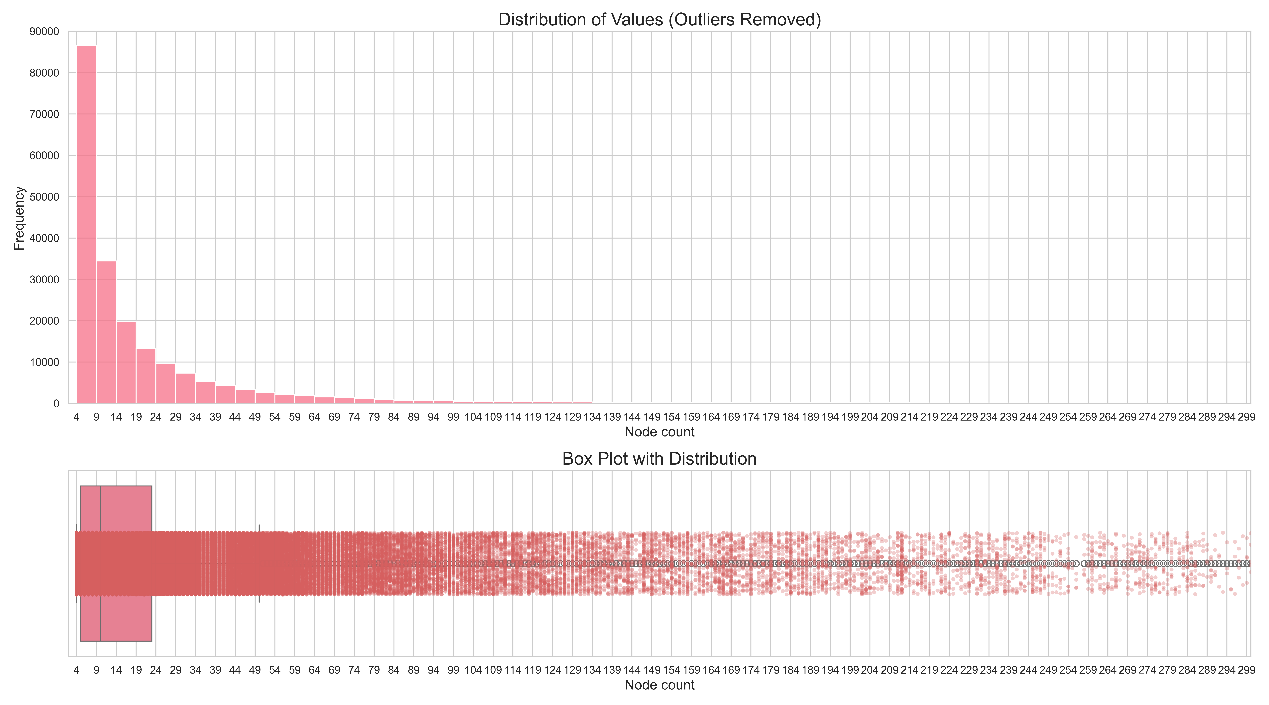

**Supplementary Fig 3.** The distribution of complex pangenome graph regions (node count >3) defined by the GRCh37 reference pangenome coordinates. The upper panel shows a histogram of node count distribution with a bin size of 4, while the lower panel presents the corresponding box plot. Statistical analysis excluded outliers above the 0.97 quantile.

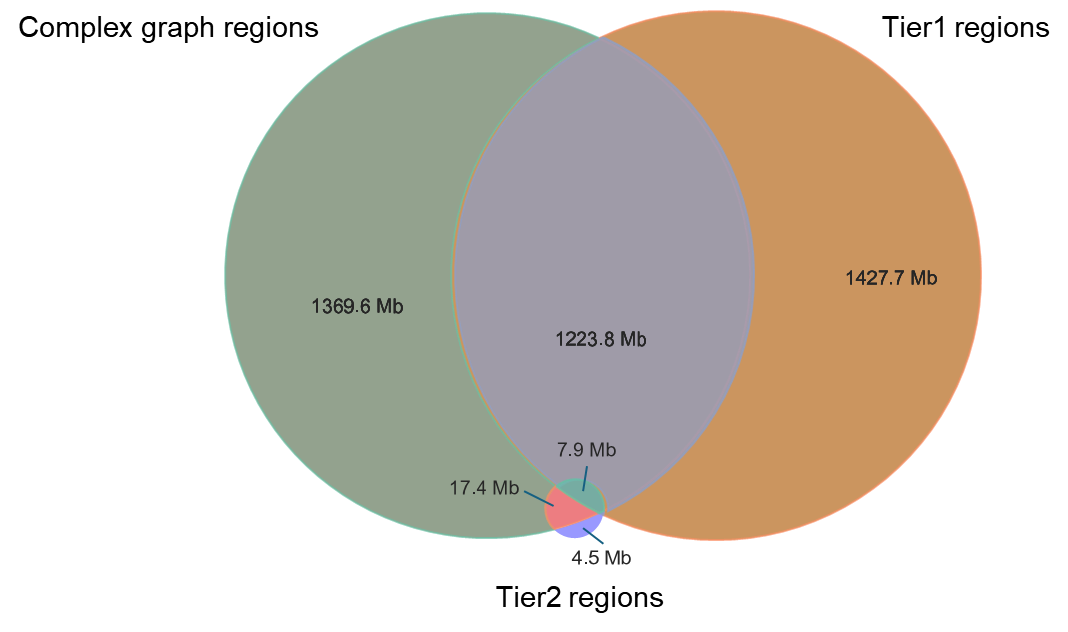

**Supplementary Fig 4**. The intersection analysis between GIAB high-confidence regions (Tier1 regions) and two types of complex regions: complex graph regions and GIAB-defined complex genomic regions (Tier2 regions). The intersection statistics were generated using bedtools intersect, with results reported in megabase pairs (Mb).

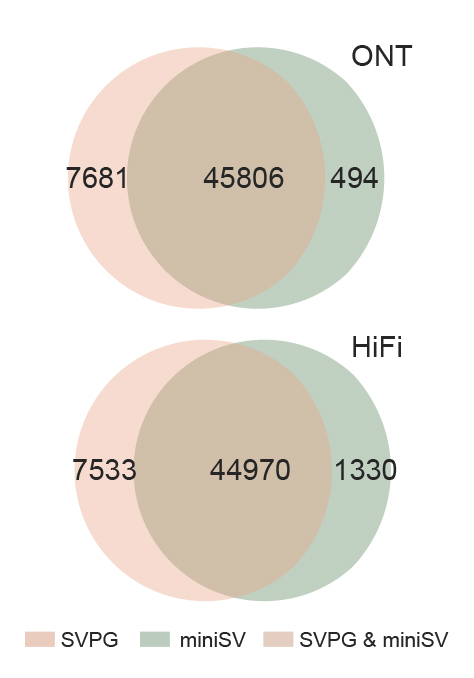

**Supplementary Fig 5.** Venn diagram of rare true positive SVs detected by SVPG and miniSV on ONT and HiFi simulated rare SVs dataset.
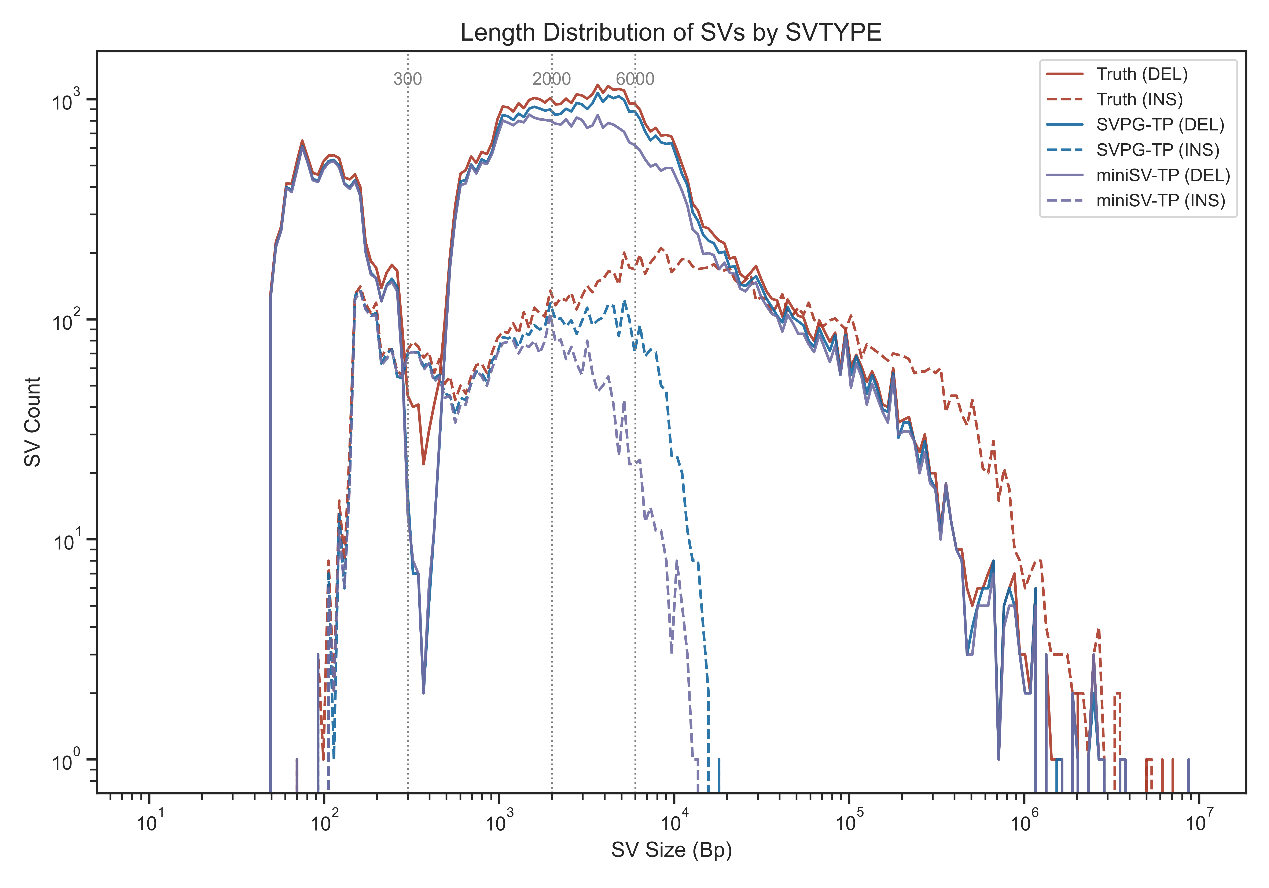

**Supplementary Fig 6.** Length distribution of rare SV truthset and true positive (TP) SV callsets from SVPG and miniSV. The TP SV callsets from Truvari evaluation result based on ONT dataset. Vertical lines at 300 bp, 2000 bp, and 6000 bp mark commonly observed SV lengths associated with mobile elements. The 300 bp and 6 kb insertions corresponded to Alu and LINE1 elements respectively, the two most abundant classes of transposable elements in the human genome (~11% and ~17% of the genome). The 2 kb SVs represent composite SVA (SINE, Variable Number Tandem Repeat, and Alu) transposons.

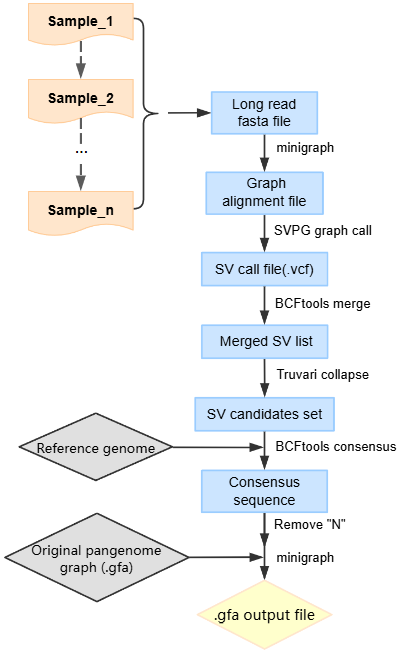

**Supplementary Fig 7.** Flowchart of the graph augmentation mode in SVPG.

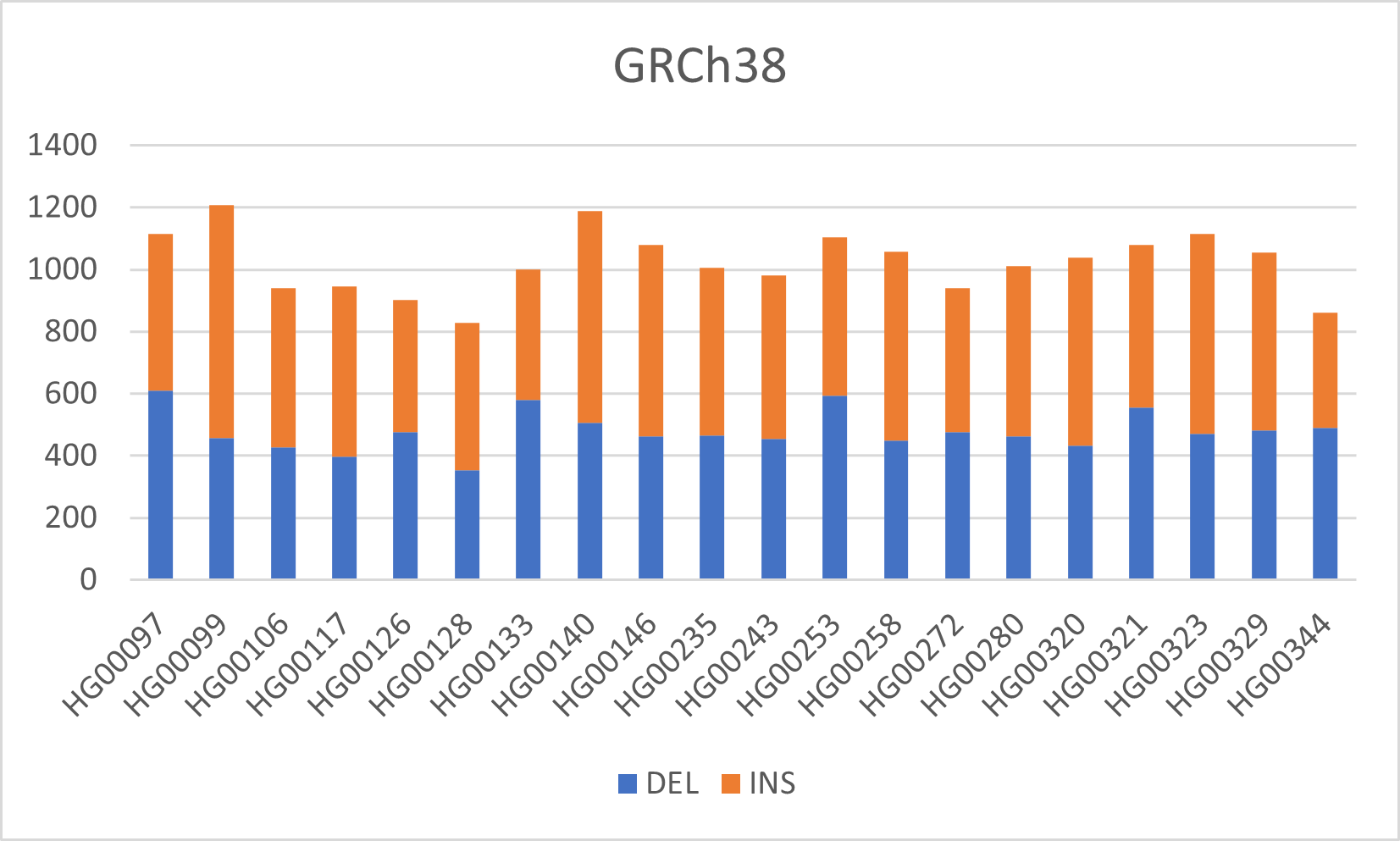

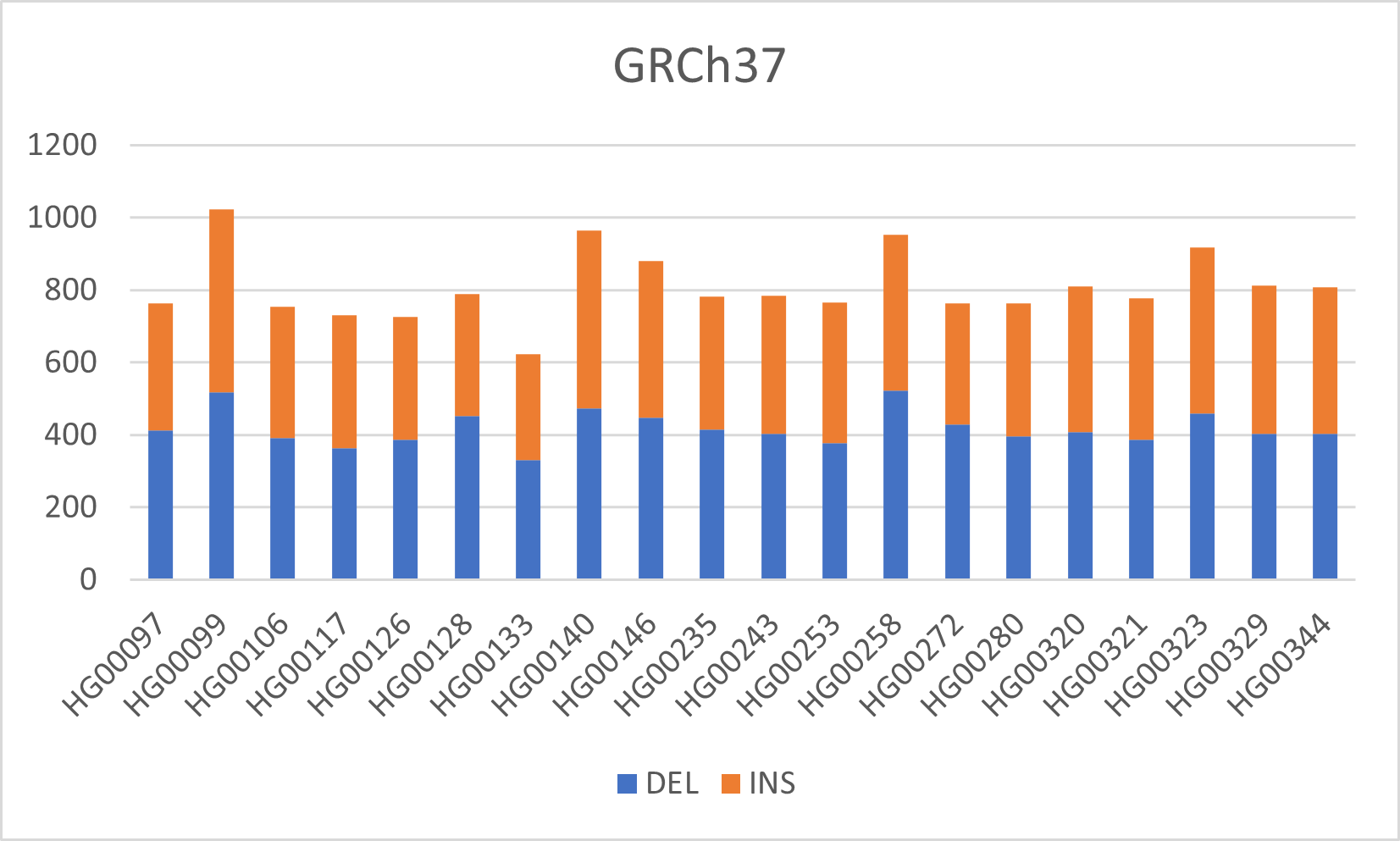

**Supplementary Fig 8.** Statistics of SV count detected by SVPG across 20 HPRC samples.

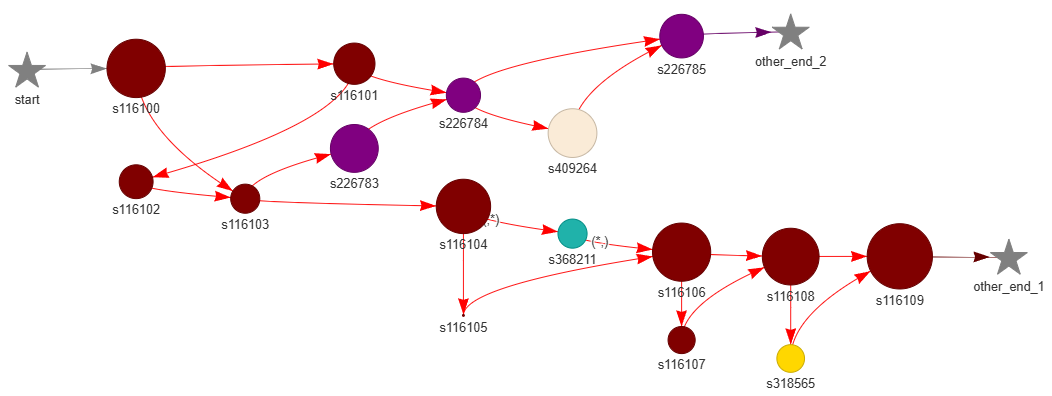
 **Supplementary Fig 9.** Local graph topology at the *MUC6* locus of hifiasm-AUG (17 nodes).

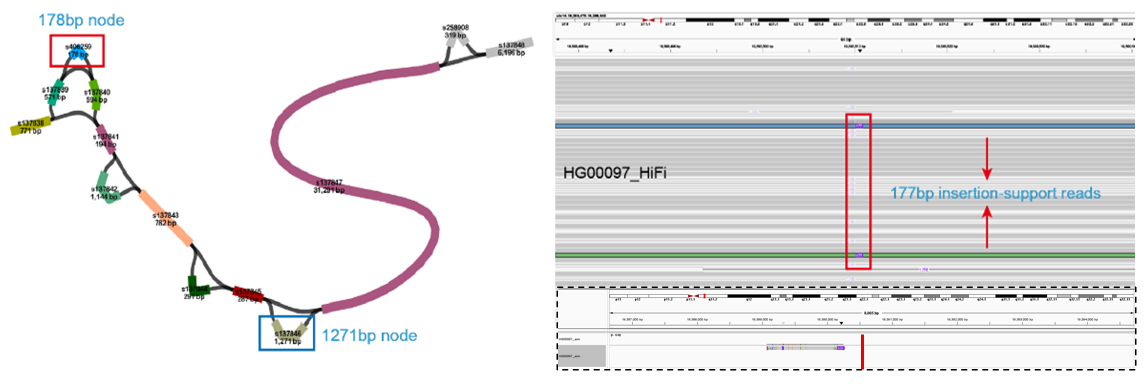

**Supplementary Fig 10.** Two SVPG-AUG-specific node examples. The Bandage view (left) shows a 178bp insertion at chr14:19,390,510 in HG00097 was identified by SVPG and represented as a bubble structure (labeled red box, node s406359). In an adjacent region, a new 1,271bp node (labeled blue box, s137847) of SVPG-AUG compared to hifiasm-AUG for the same region. By examining the VCF files of 20 new samples called by SVPG, we confirmed this variant originated from the HG00117 sample (chr14:19,393,631-19,394,902). The IGV view (top right) shows read support for the 178bp insertion in HG00097 at the chr14:19,390,510 position, whereas the bottom panel indicates a local assembly failure by hifiasm in this region.

**Supplementary Notes**

**Supplementary Note 1. Validation of the five *de novo* SVs of SVPG and the 13 *de novo* SVs of Sniffles2**

Validation of discordant SVs called by SVPG and Sniffles2 in HG002 sample across chromosomes 1-3, using IGV screenshots and GIAB benchmark set (high-confidence regions). Each IGV snapshot spans the region covering the start and end positions of a *de novo* SV called in HG002, extended by 1000 bp on both sides. At the top of the snapshot, the nearest SV record from the GIAB benchmark set is shown; if no record is found within 1000 bp of the *de novo* SV, it is labeled as “NA”. On the left side of the snapshot, SVs called by different tools are displayed for HG002, HG003, and HG004 within the same extended region. For HG003 and HG004, only the closest SV to the *de novo* SV position is displayed; if this SV is more than 1000 bp away, it is also labeled as “NA”.

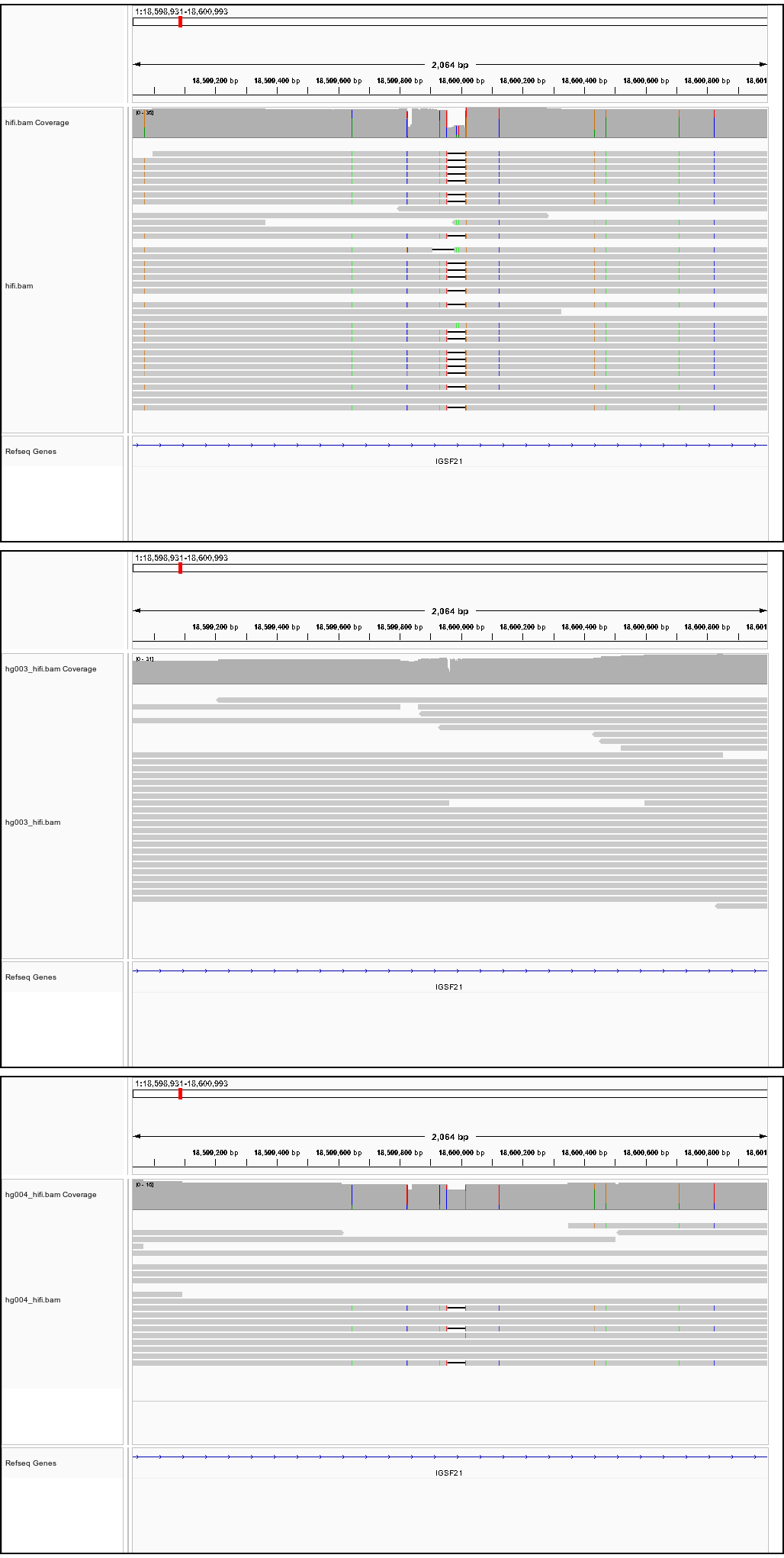

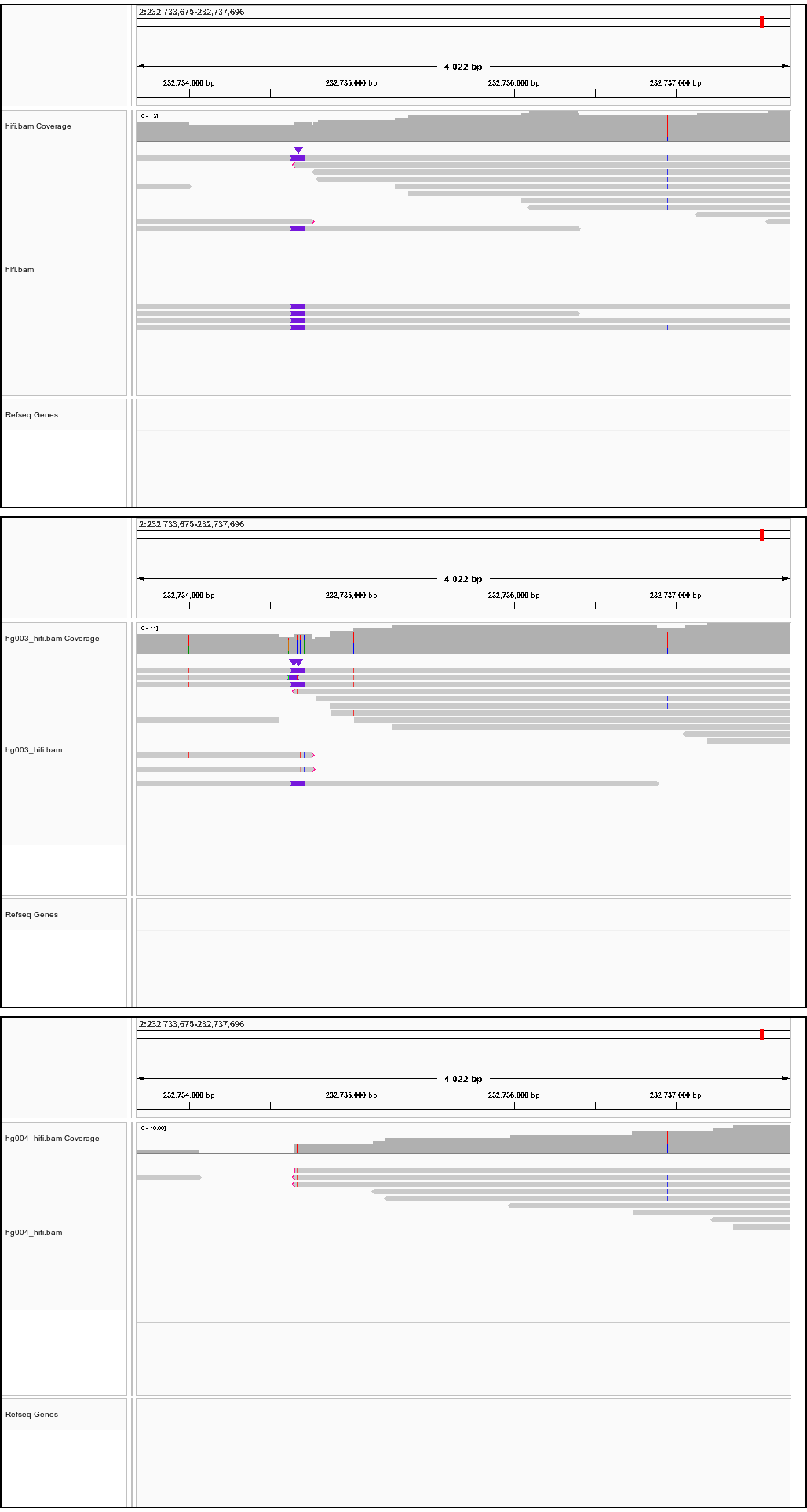

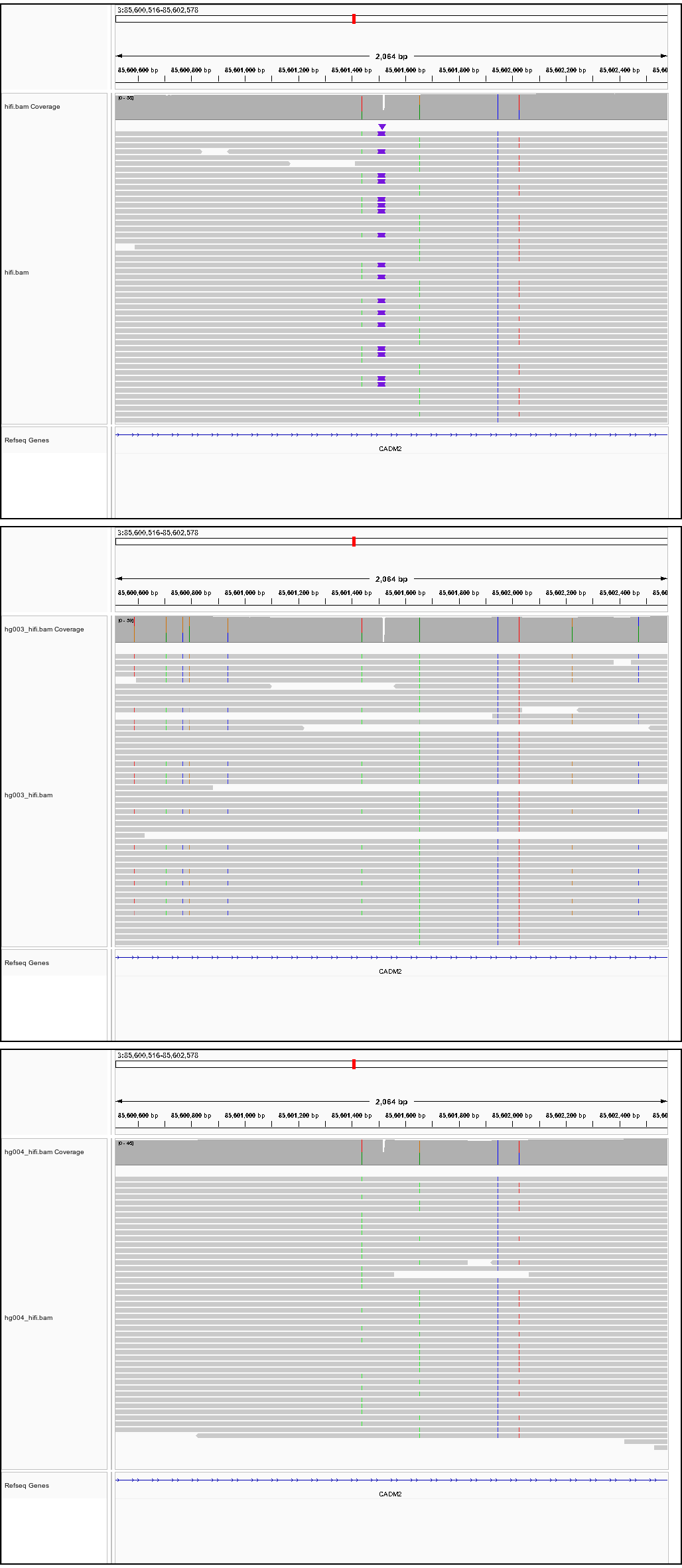

Successfully verified true positive *de novo* SV SVs

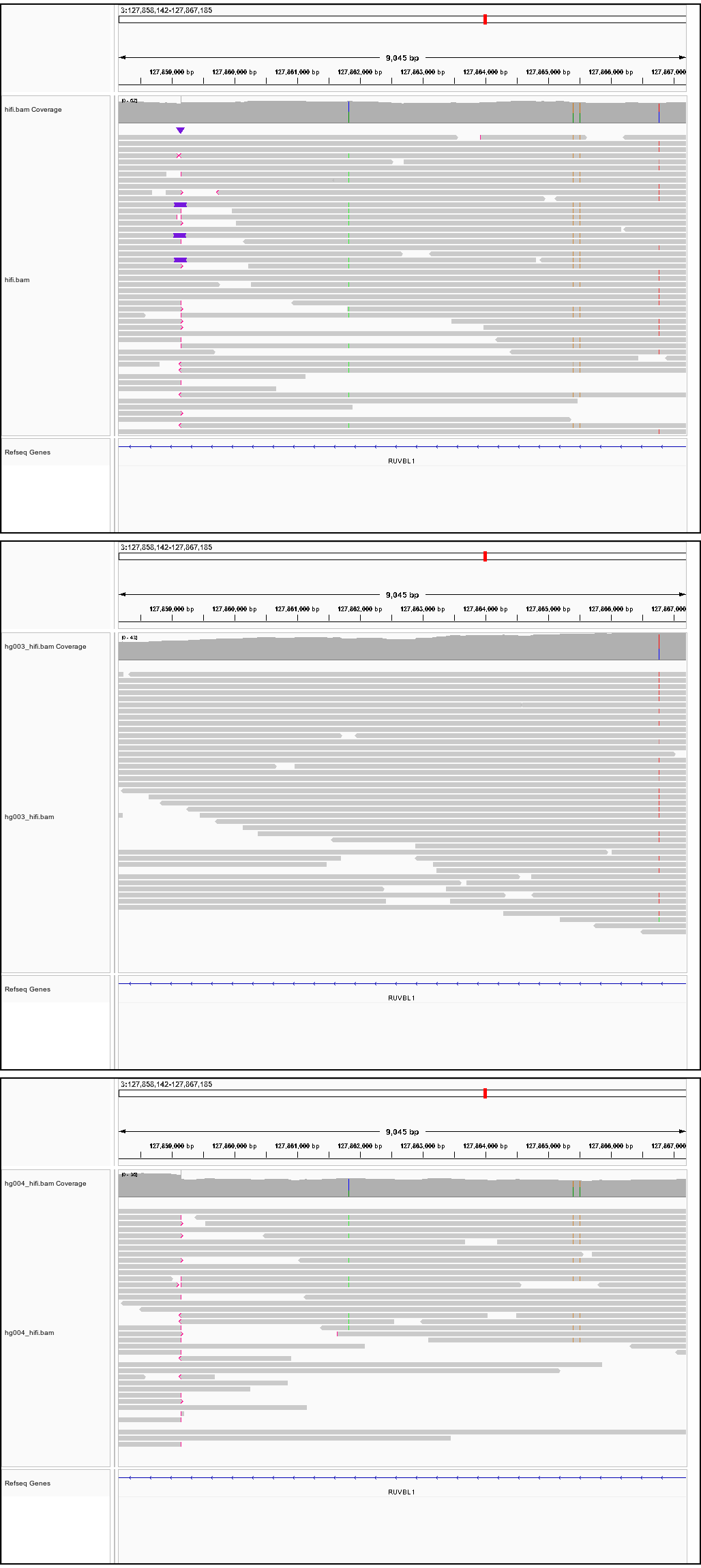

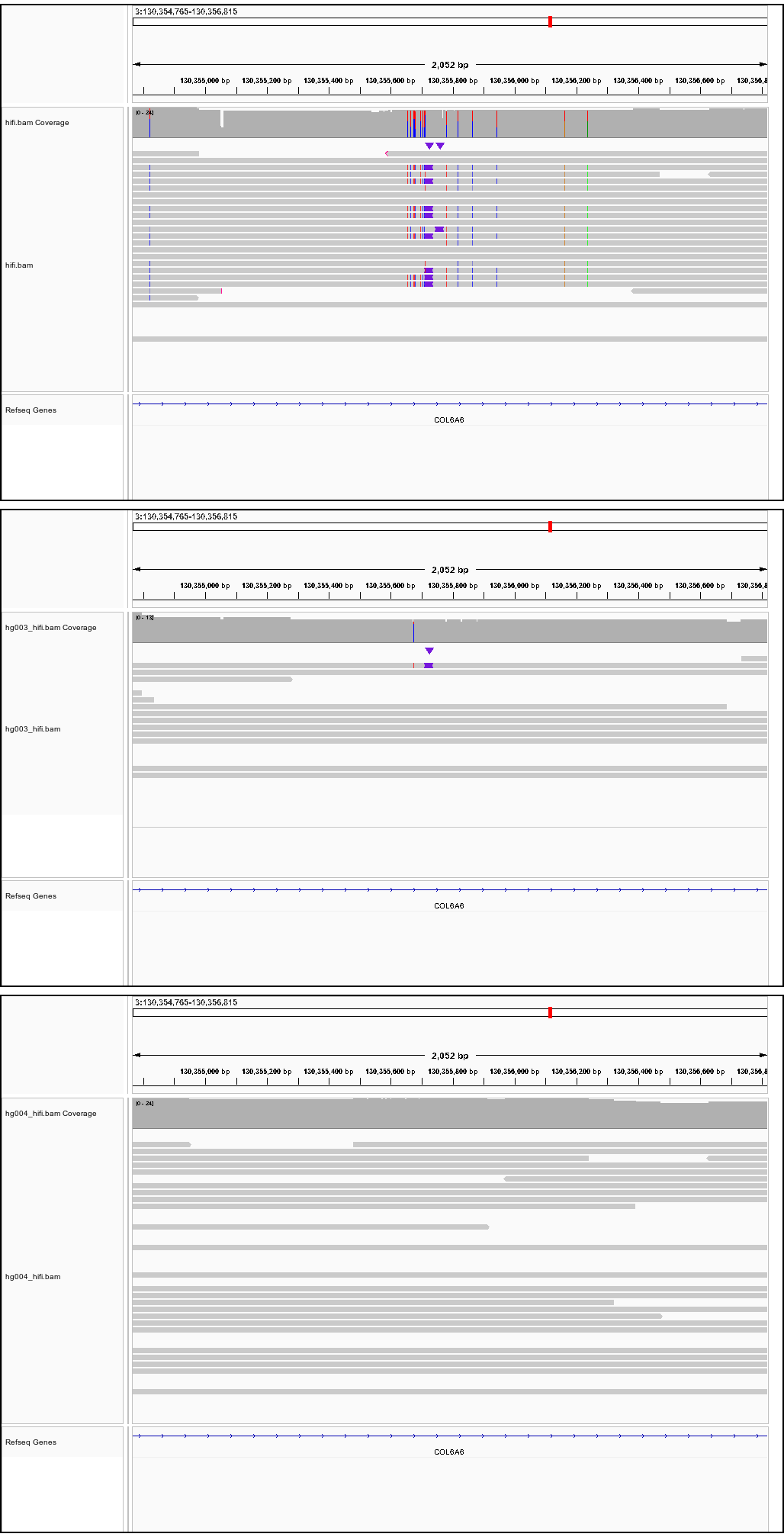

Successfully verified true positive *de novo* SV SVs

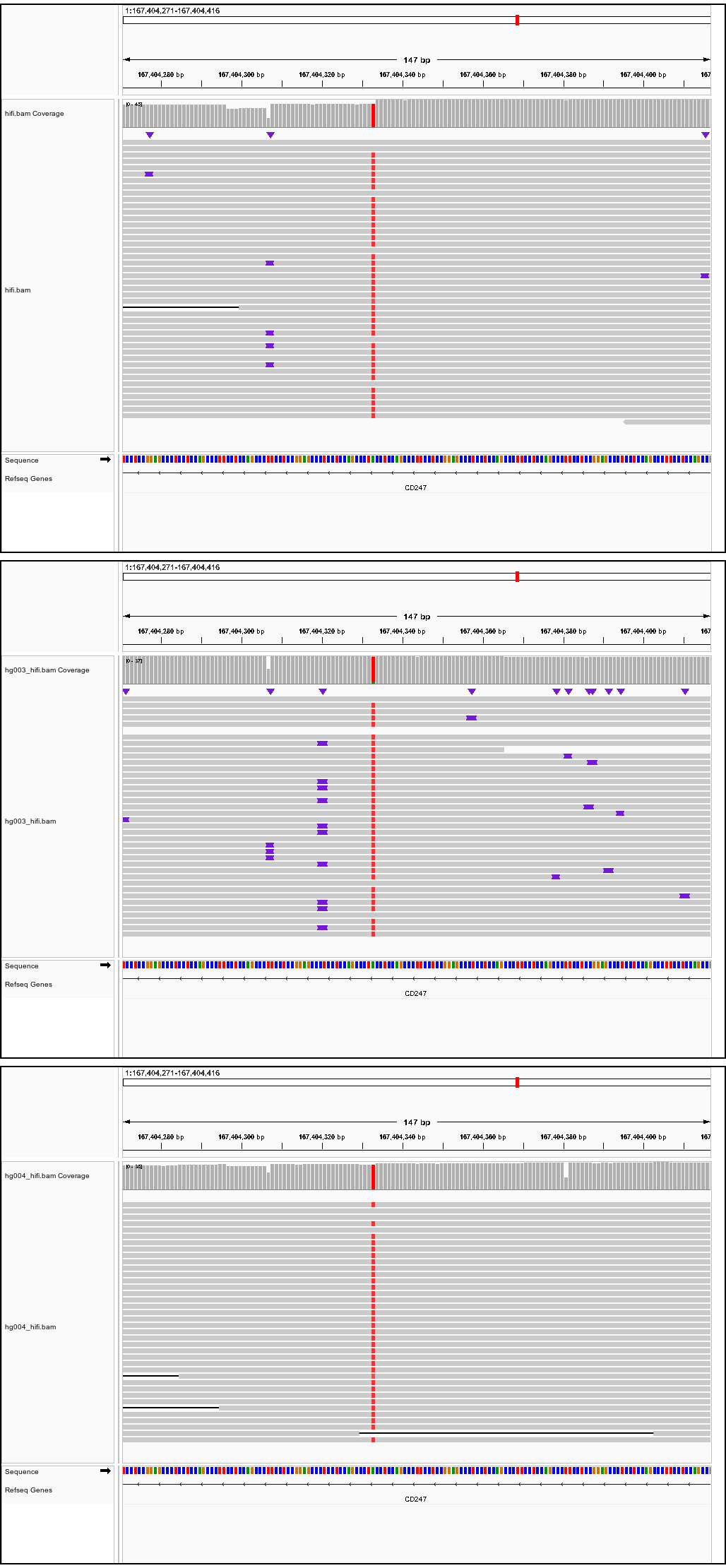

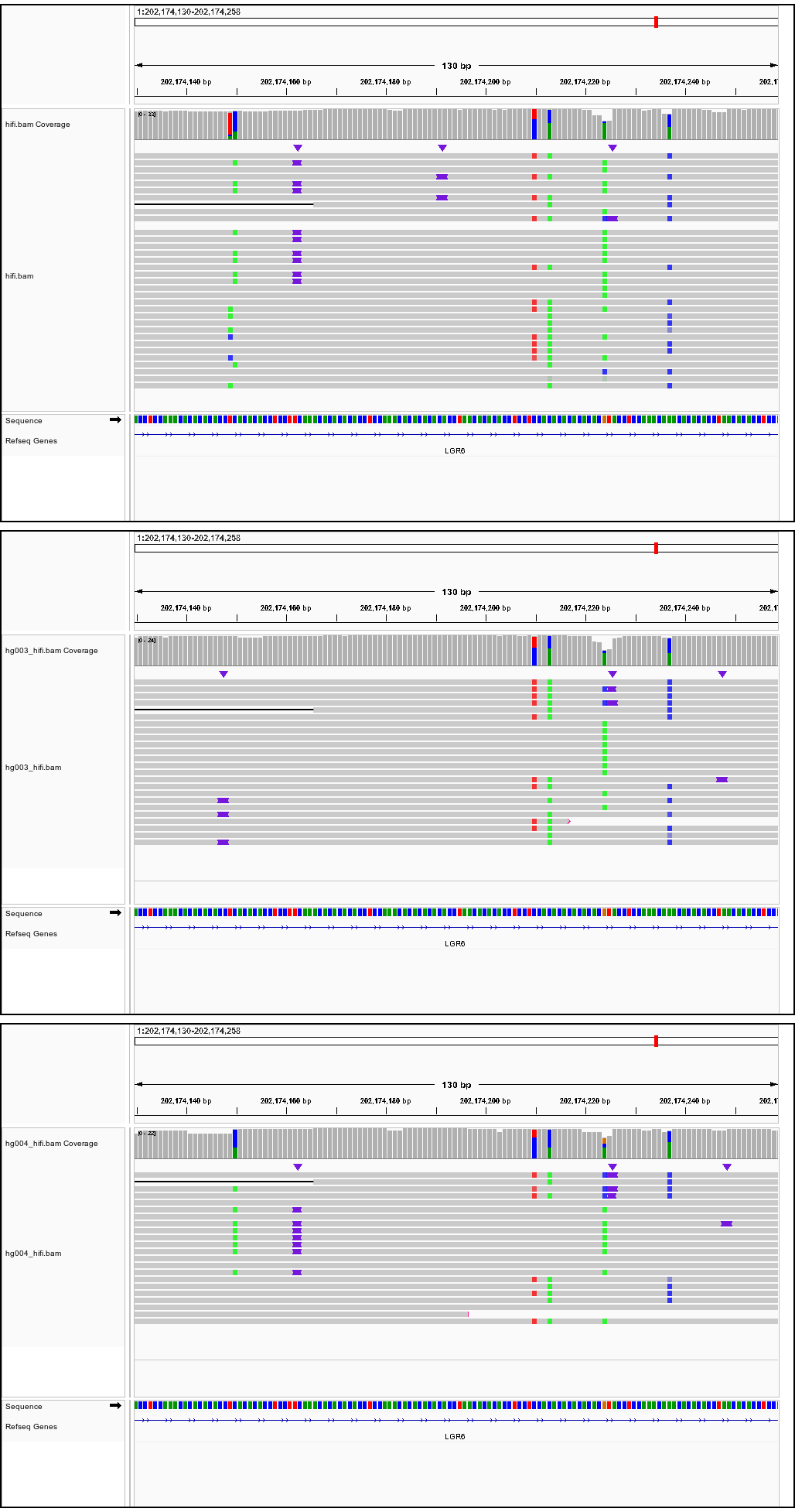

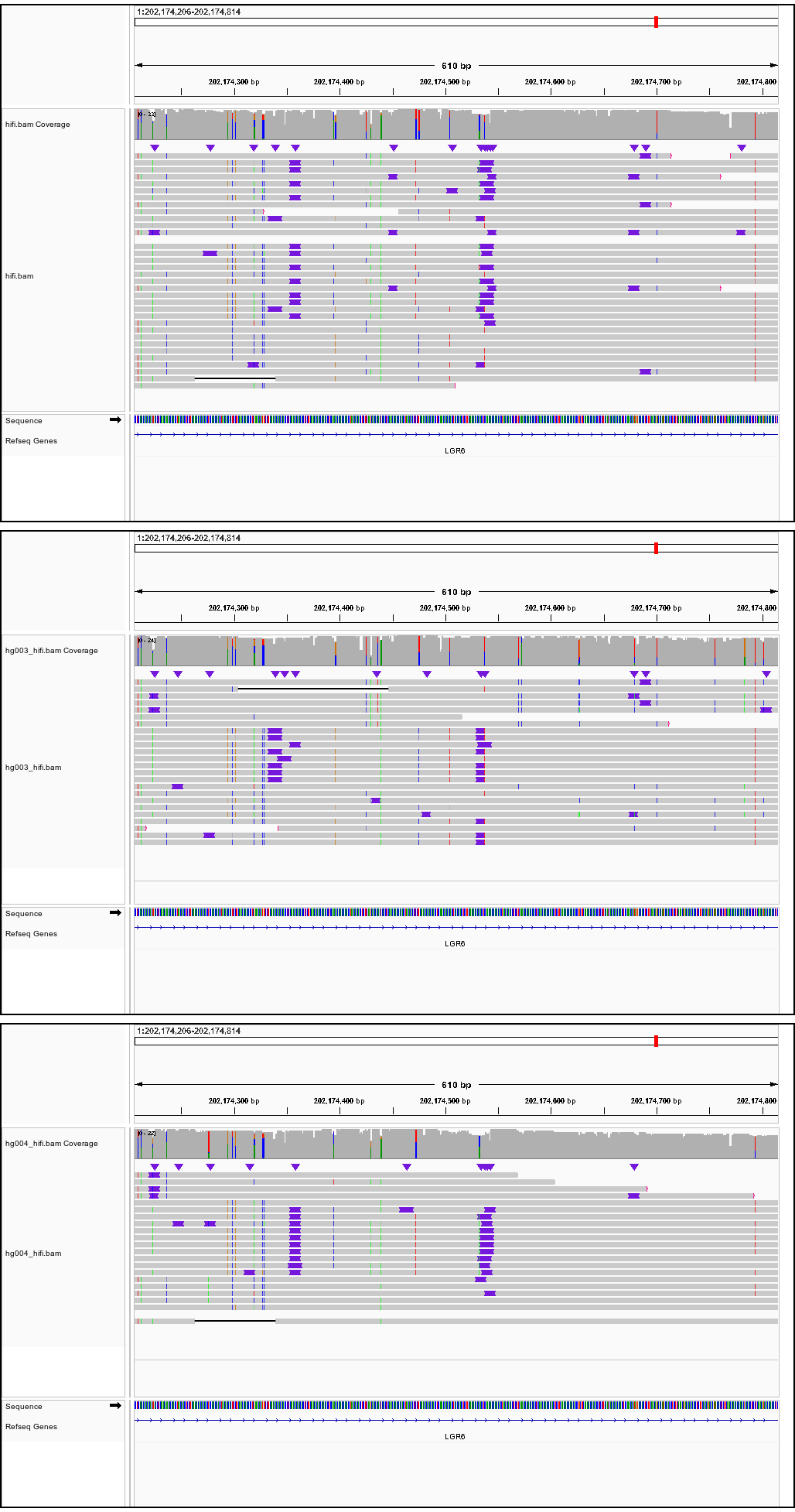

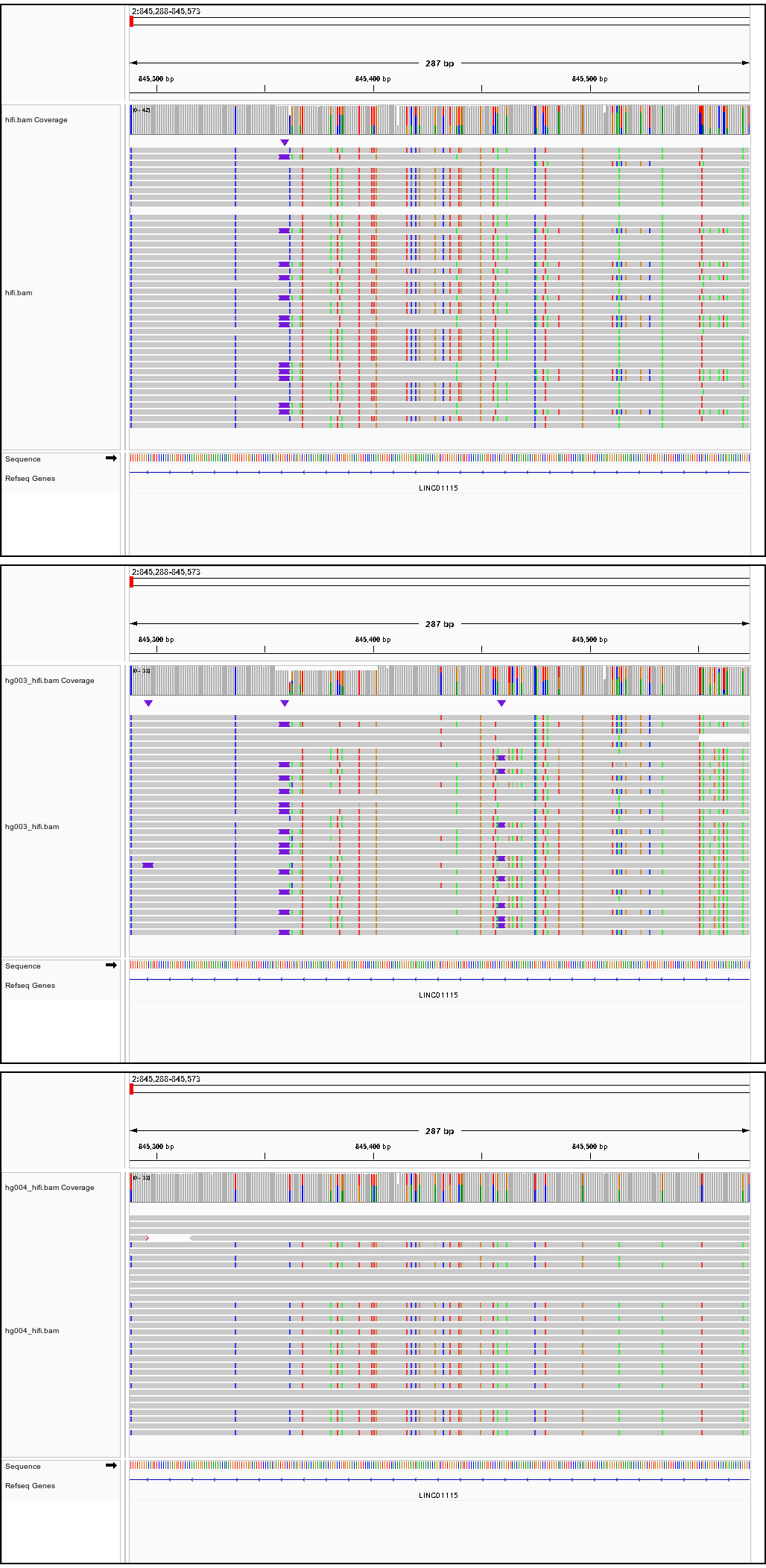

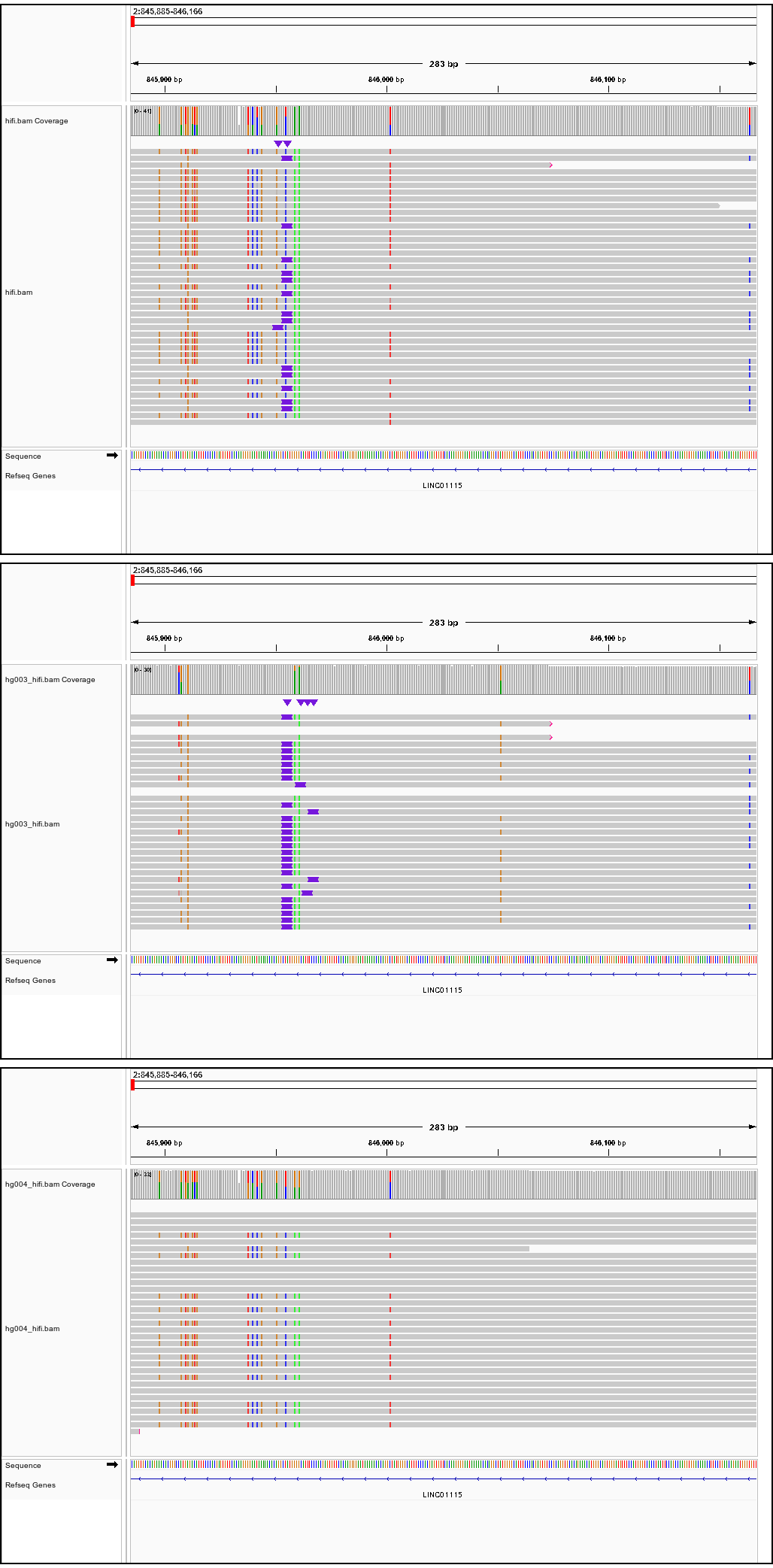

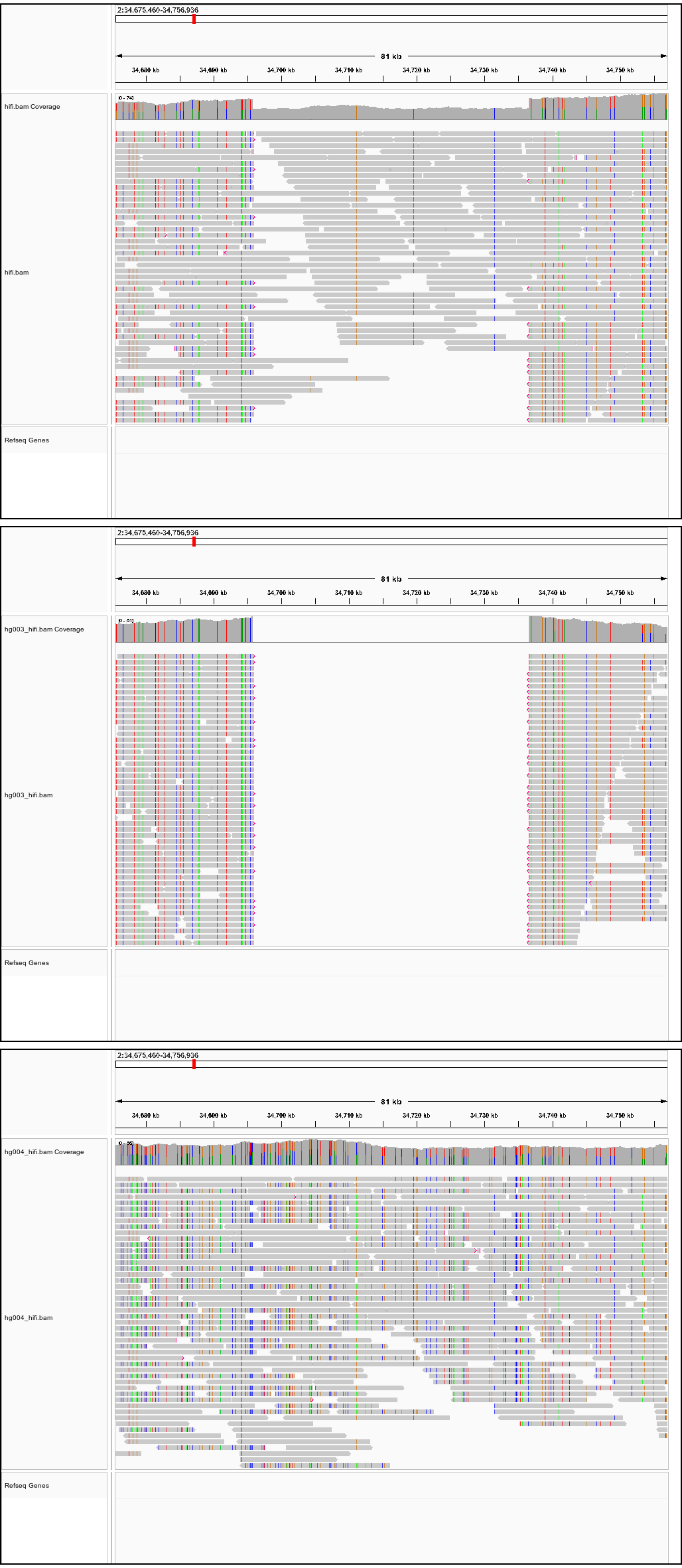

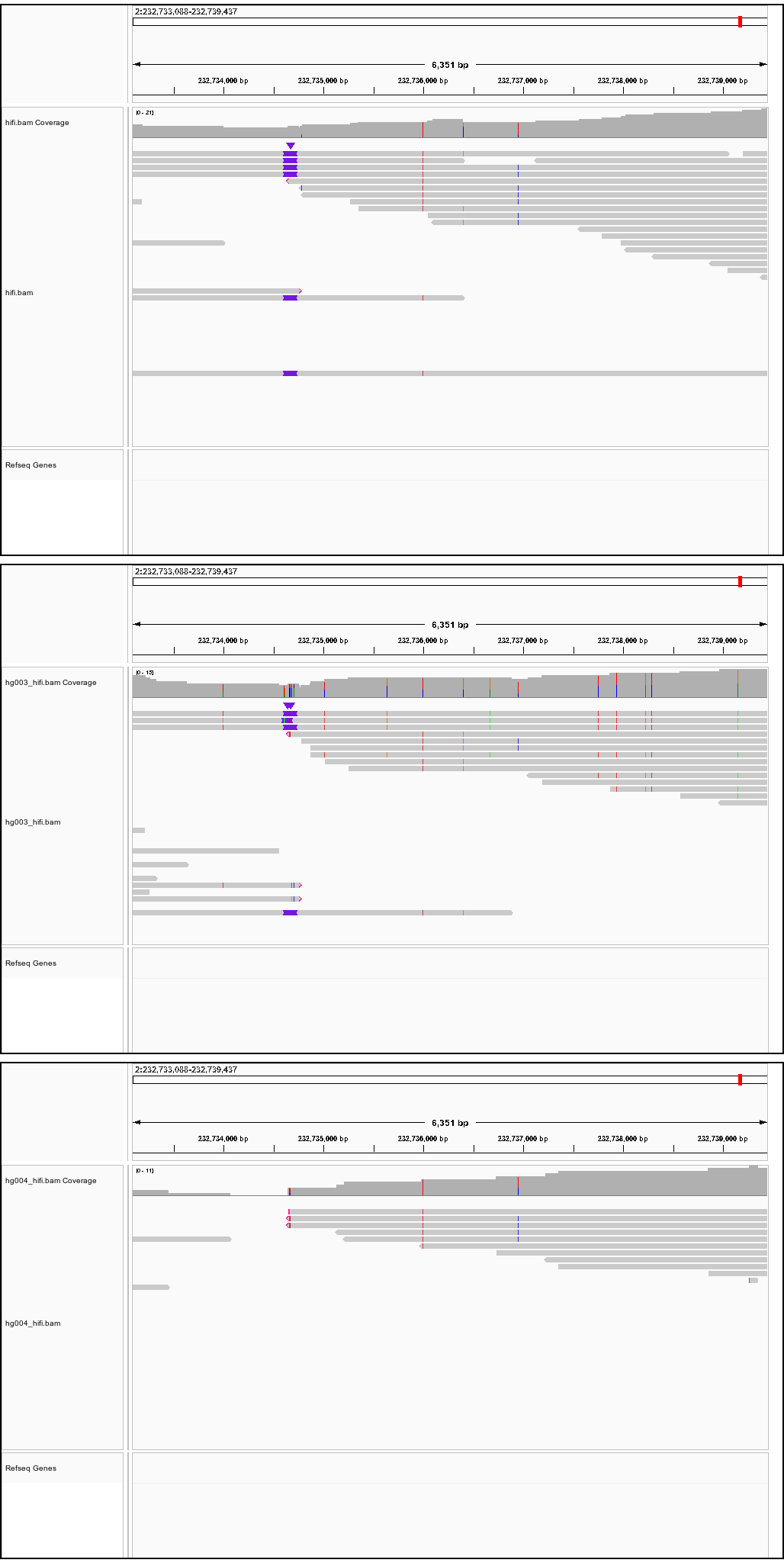

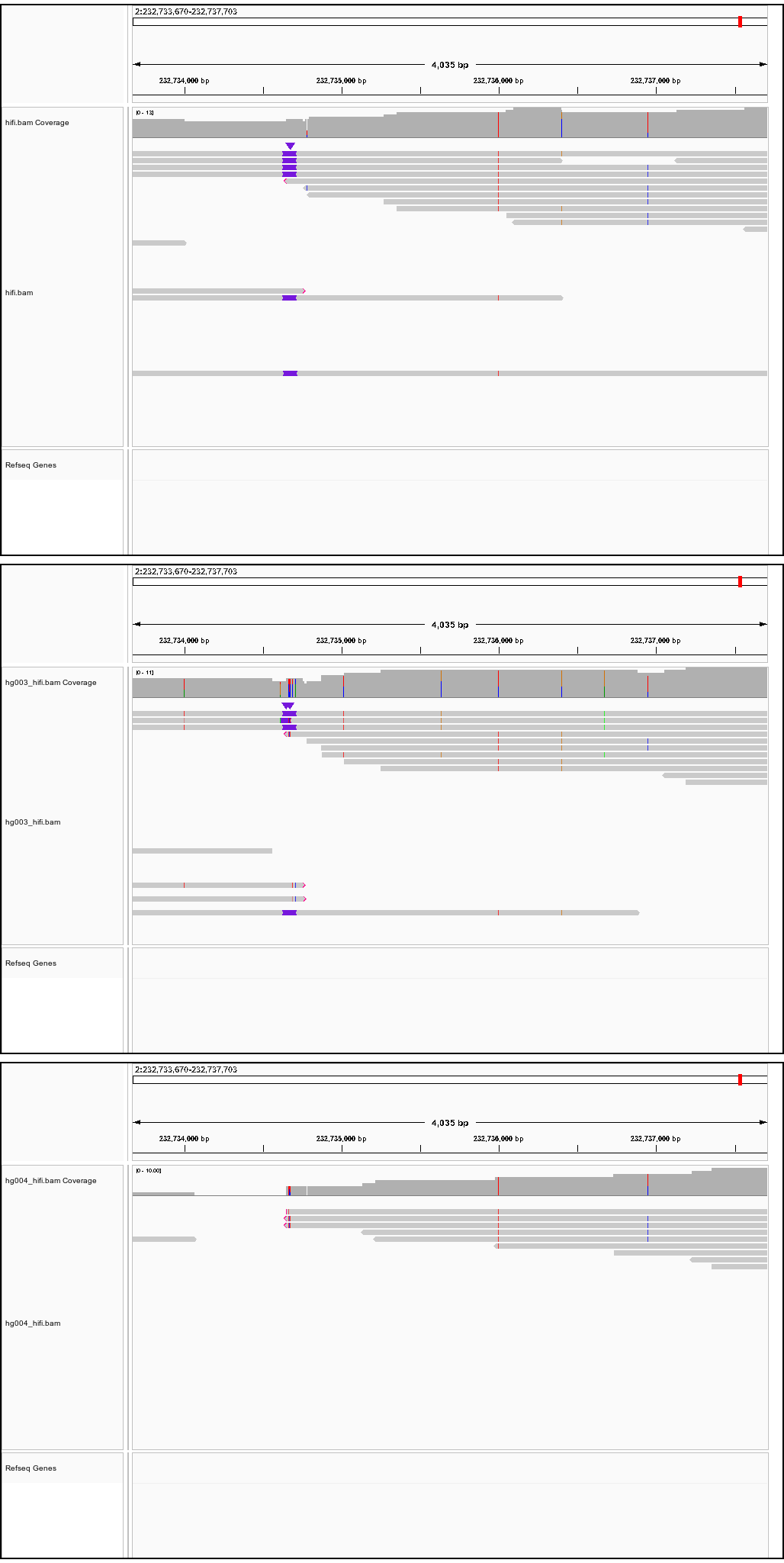

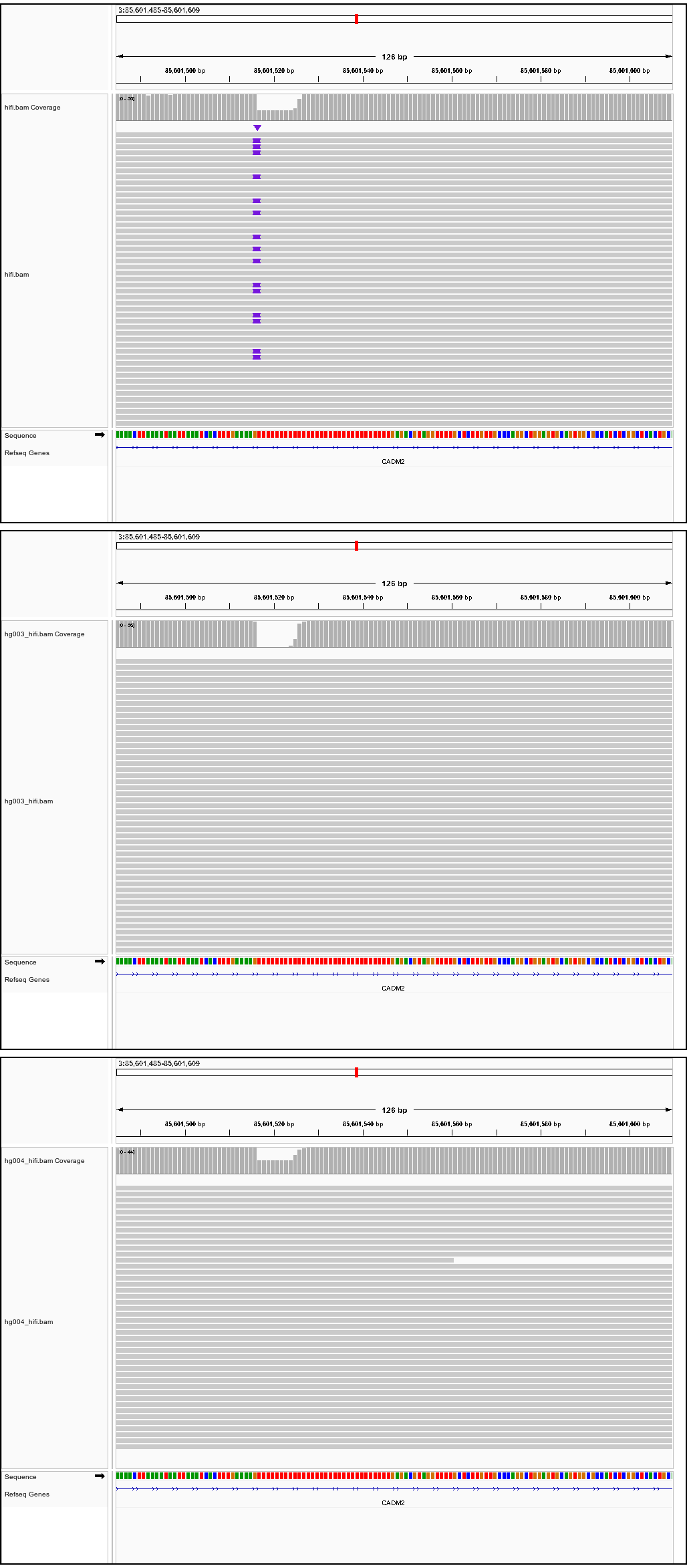

Successfully verified true positive *de novo SV* SVs

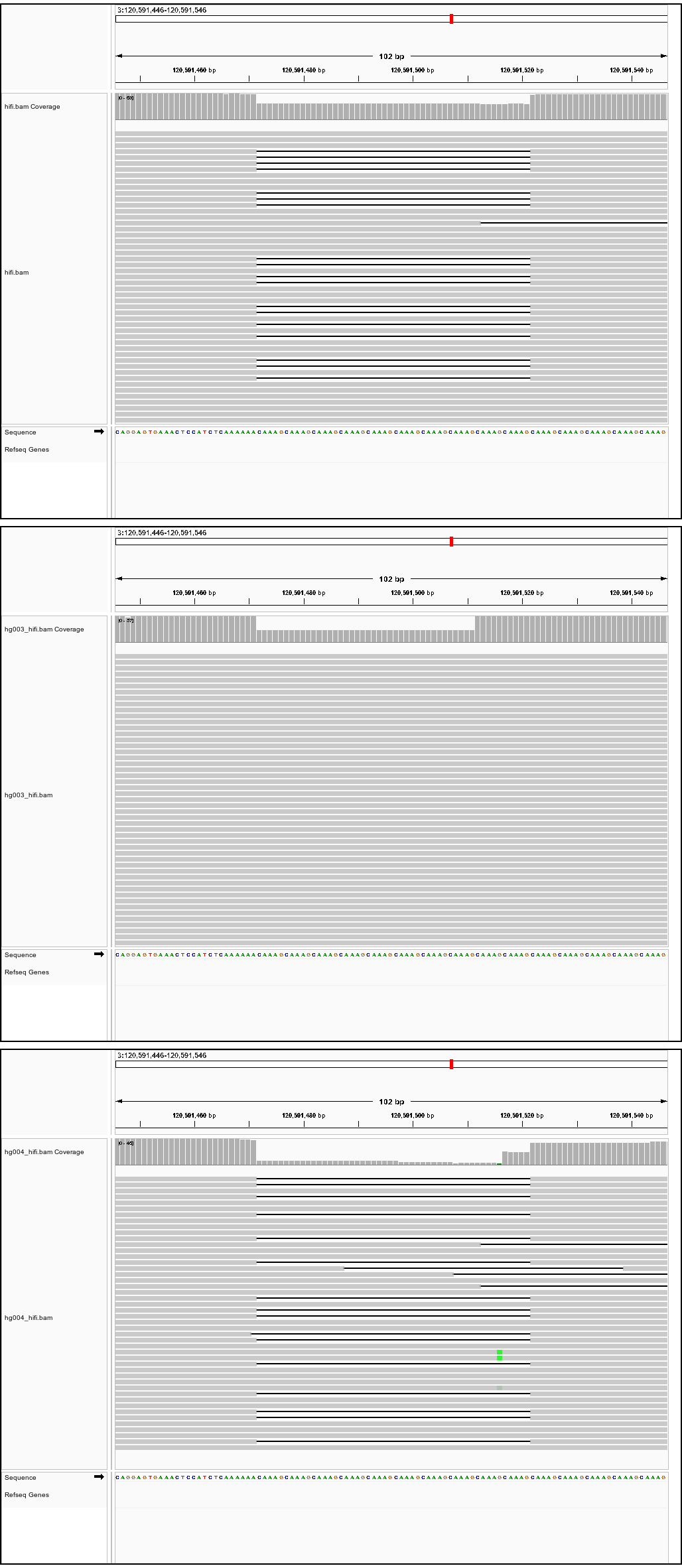

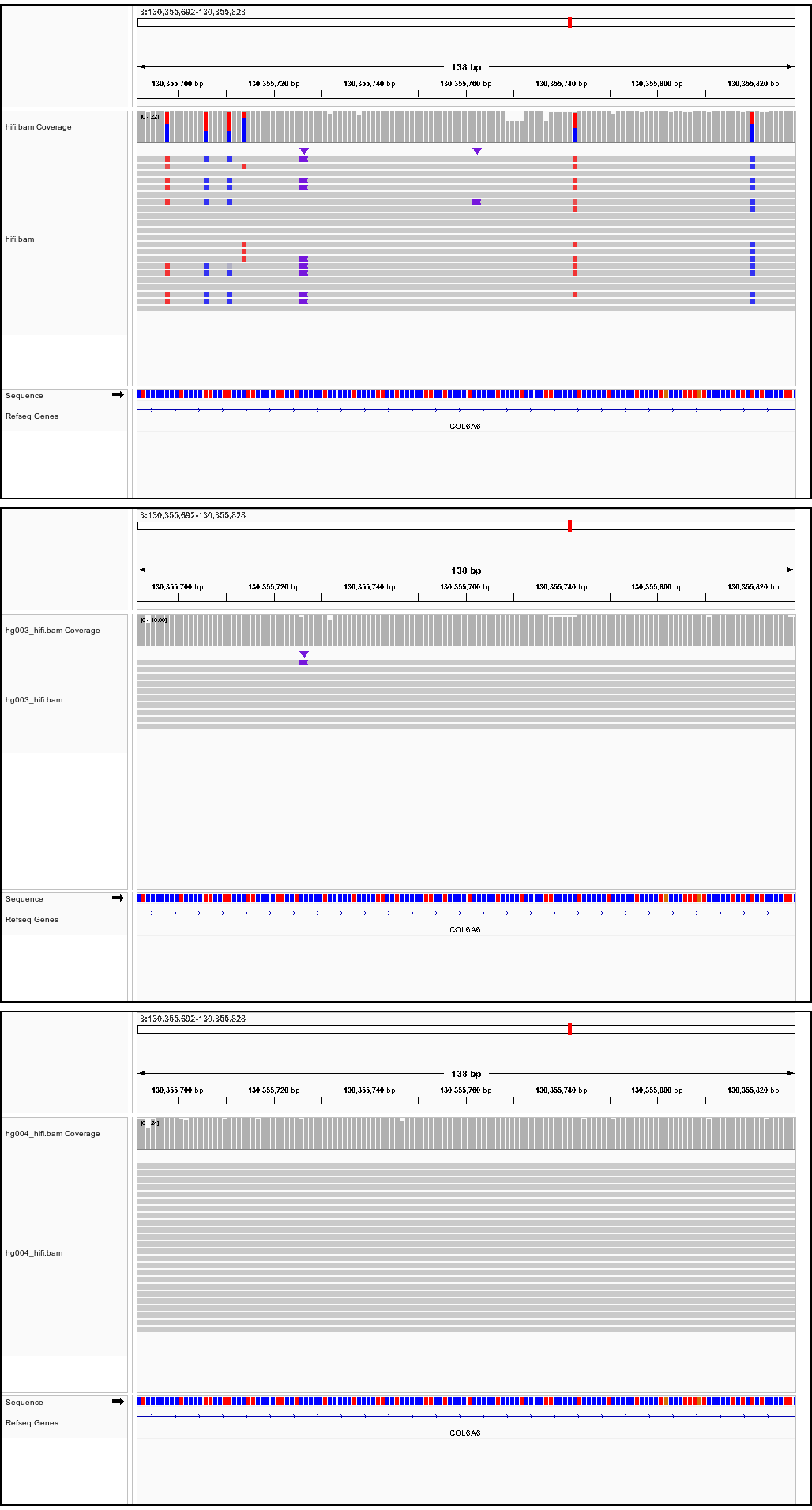

Successfully verified true positive *de novo* SV SVs

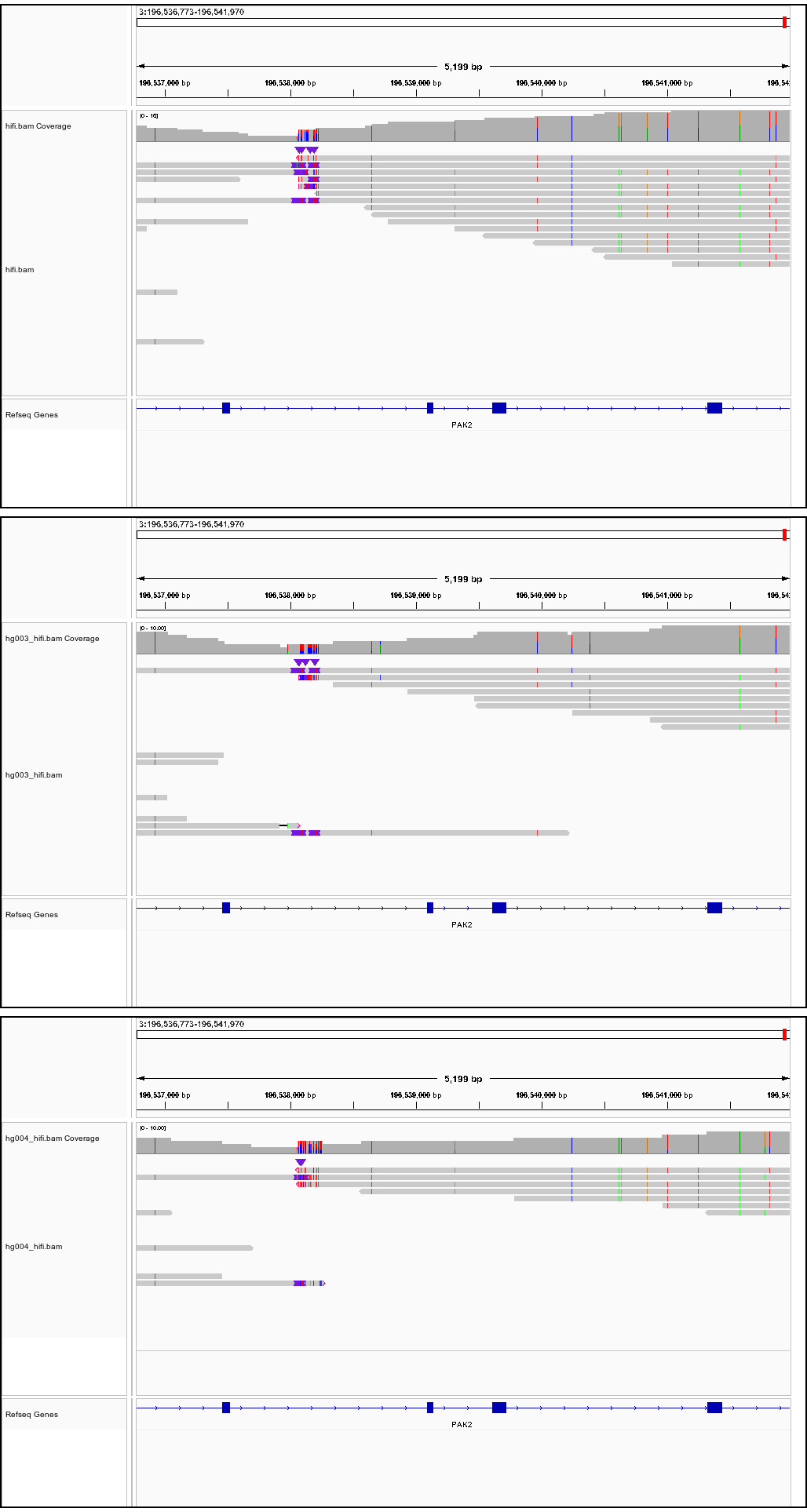

**Supplementary Note 2. Validation of four SVPG-missed SVs in HG008 against the GIAB benchmark set.**

Validation of seven SVs missed by SVPG based on the GIAB benchmark set, four of which are represented in the pangenome graph. The IGV screenshots support the credibility of the benchmark calls, while the Bandage visualizations illustrate the pangenome context in the corresponding regions (Supplementary Table 9 for overlap information detail).

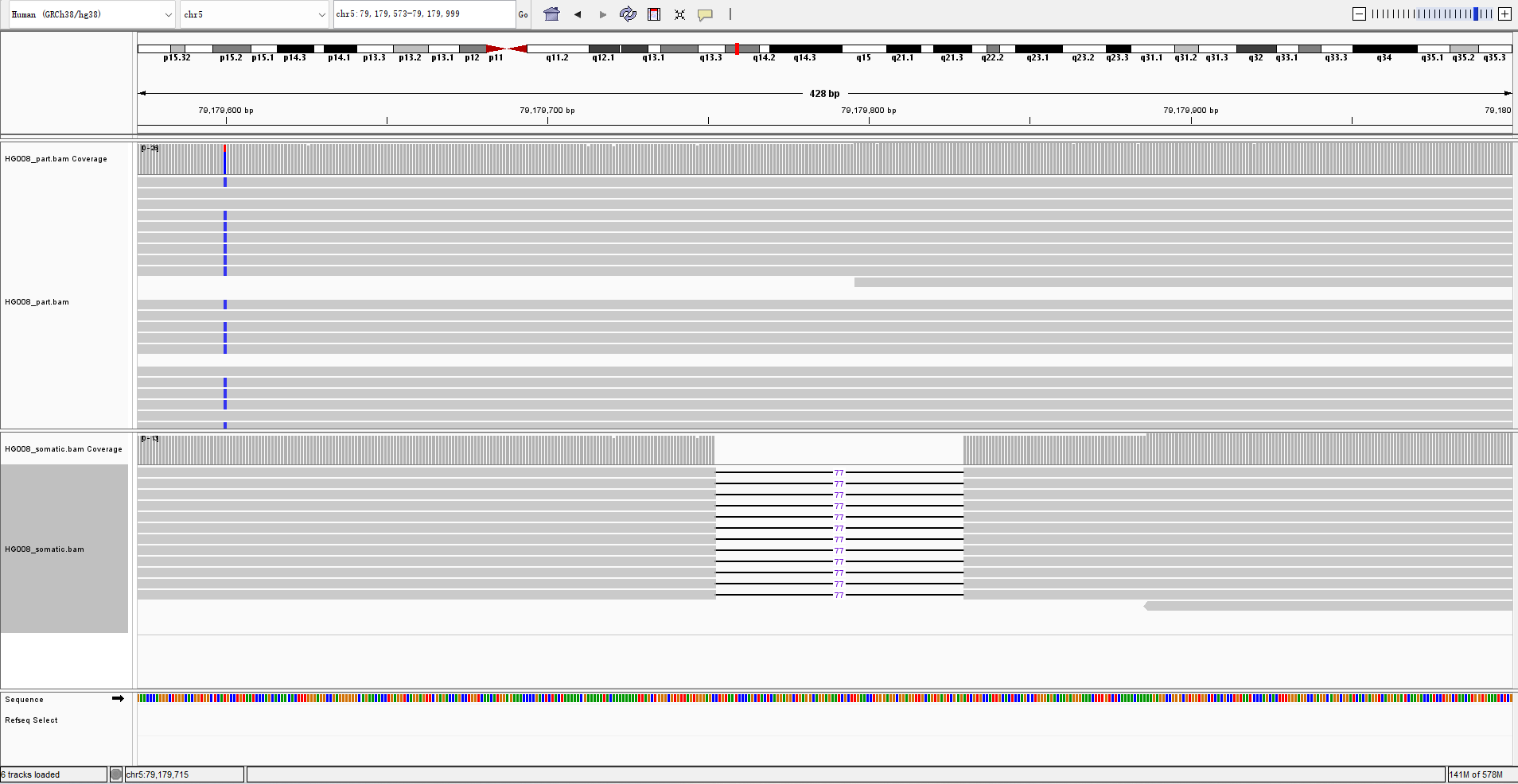

**Chr5:** **79179752-79179829, DEL**

**Control**

**Tumor**

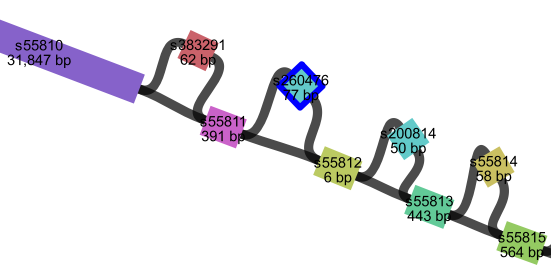

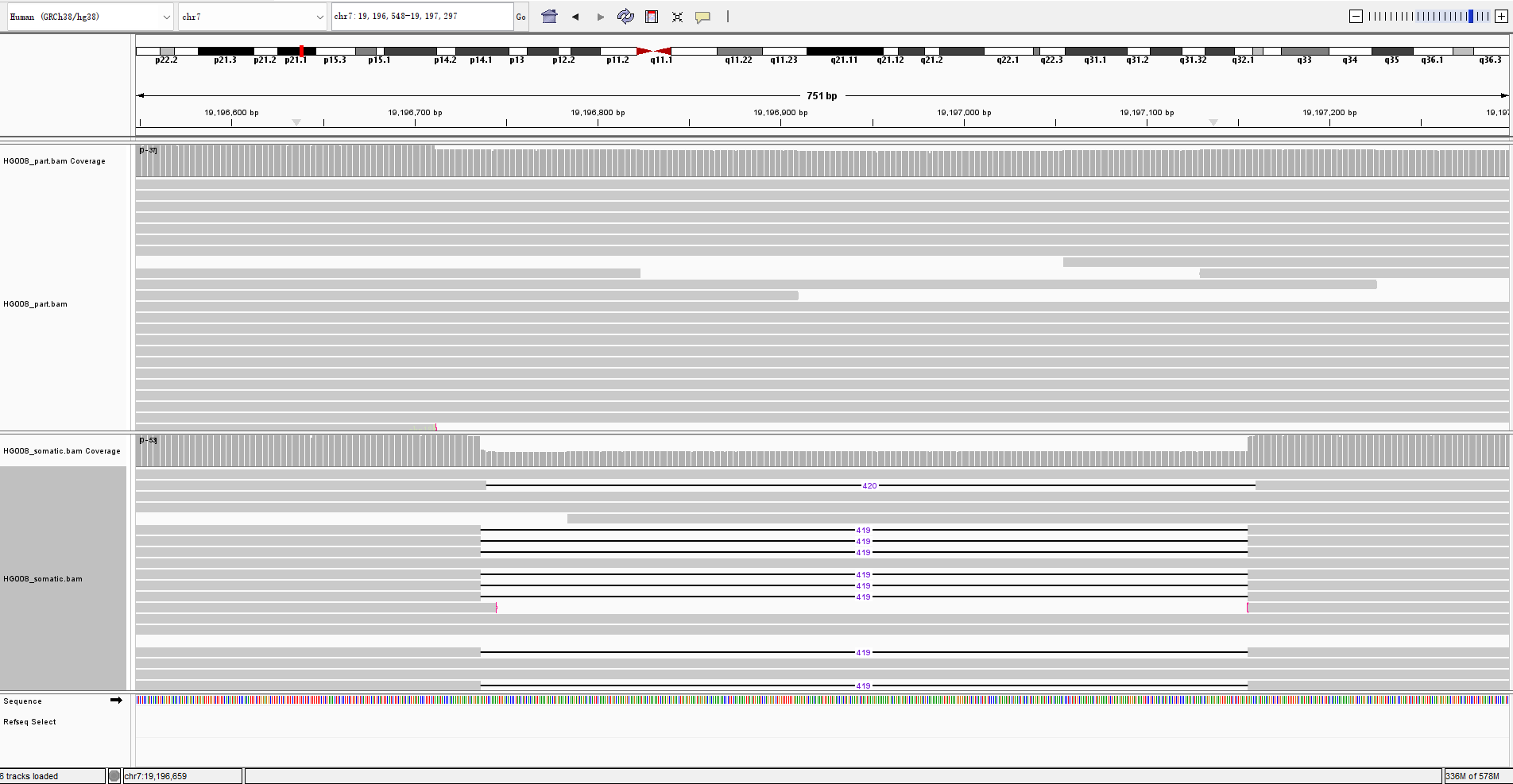

**Chr7:** **19196736-19197155, DEL**

**Control**

**Tumor**

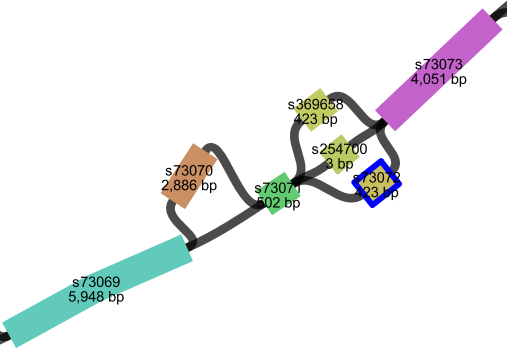

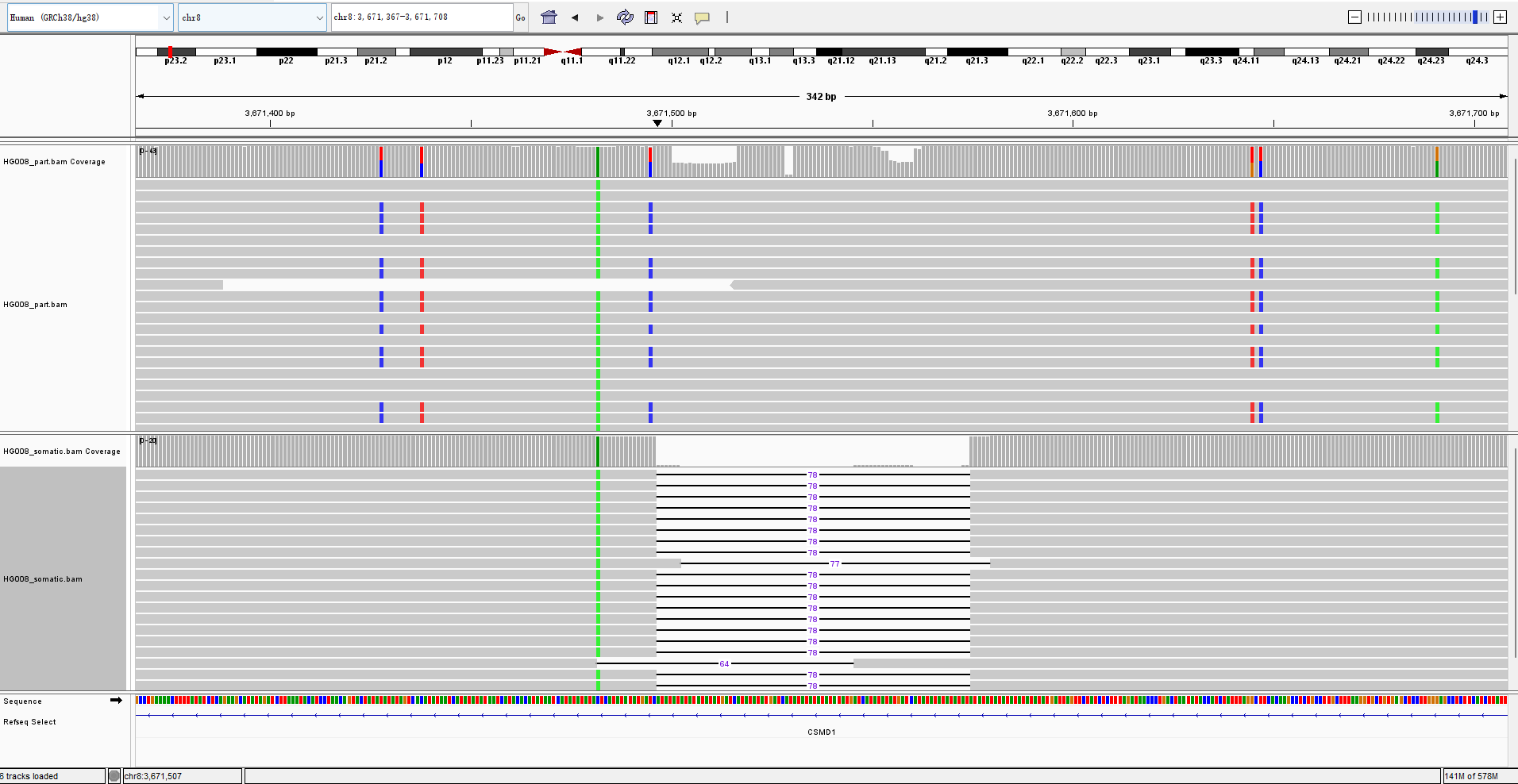

**Control**

**Tumor**

**Chr8:** **3671496-3671574, DEL**

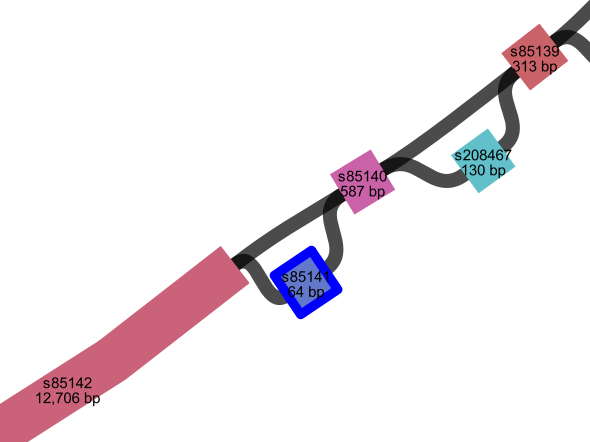

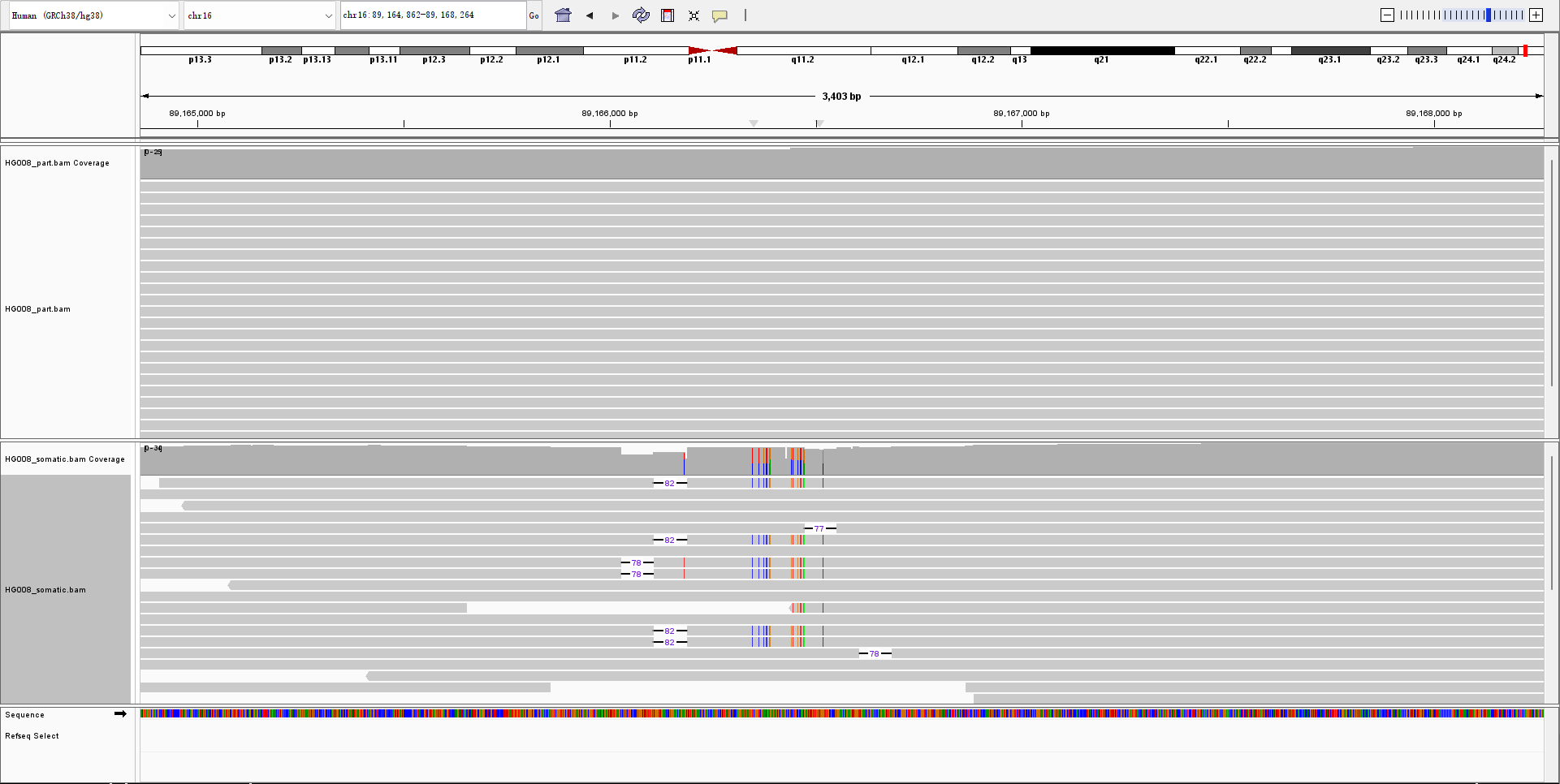

**Tumor**

**Control**

**Chr13:** **66842467-73136072, DEL**

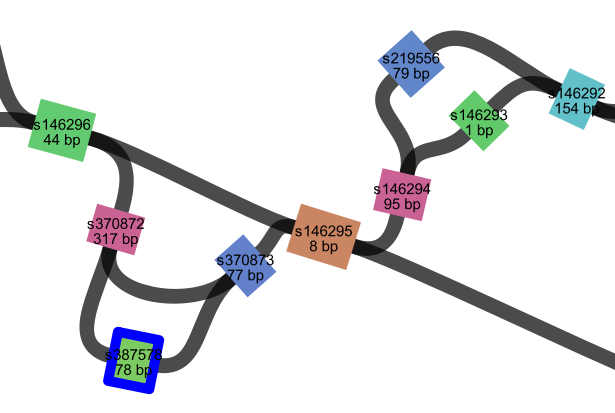

**Supplementary Note 3. Commands used**

1	Command lines for each caller

**SVPG (v1.3)**

Pangenome-guided call mode:

*SVPG call --working_dir /path/to/output --bam /path/to/sample.bam --ref /path/to/ref.fa --gfa /path/to/pangenome.gfa -s {min_support} -t 16*

Pangenome-based mode (graph call mode):

*SVPG graph-call --working_dir /path/to/output --ref /path/to/ref.fa --gaf /path/to/sample.gaf --gfa /path/to/pangenome.gfa -s {min_support} -t 16*

Graph augment mode:

*SVPG augment --working_dir /path/to/samples --ref /path/to/ref.fa --gfa /path/to/pangenome.gfa*

**Sniffles2 (v2.3.3)**

*sniffles --input /path/to/sample.bam --vcf /path/to/output.vcf -t 16*

**cuteSV (v2.1.1)**

HiFi:

*cuteSV /path/to/sample.bam /path/to/ref.fa /path/to/output/output.vcf /path/to/output -s {min-support} --genotype -t 16 --max_cluster_bias_INS 1000 --diff_ratio_merging_INS 0.9 --max_cluster_bias_DEL 1000 --diff_ratio_merging_DEL 0.5*

ONT:

*cuteSV /path/to/sample.bam /path/to/ref.fa /path/to/output/output.vcf /path/to/output -s {min-support} --genotype -t 16 --max_cluster_bias_INS 100 --diff_ratio_merging_INS 0.3 max_cluster_bias_DEL 100 --diff_ratio_merging_DEL 0.3*

**SVIM (v1.4.1)**

*svim alignment /path/to/output/variants.vcf /path/to/sample.bam /path/to/ref.fa*

**DeBreak (v1.2)**

*Debreak --bam /path/to/sample.bam --outpath /path/to/output --ref /path/to/ref.fa --poa --rescue_large_ins --rescue_dup -t 16*

**Sawfish (v2.0.3)**

Discover step:

*sawfish discover --threads 16 --ref /path/to/ref.fa --bam /path/to/sample.bam --output-dir /path/to/discover_dir*

Joint-call step:

*sawfish joint-call --threads 16 --sample /path/to/discover_dir --output-dir /path/to/output_call*

**miniSV (r29)**

*minisv.js extract -l 50 -b data/hs38.cen-mask.bed /path/to/sample.gaf > /path/to/variants.rsv;*

*cat /path/to/variants.rsv | sort -k1,1 -k2,2n -S4g | minisv.js merge -c {min_support} - >/path/to/variants.msv;*

*minisv.js genvcf /path/to/variants.msv > /path/to/ variants.vcf*

**SVarp (v1.0)**

*build/svarp -a /path/to/sample.gaf -g /path/to/pangenome.gfa --fasta /path/to/sample.fasta -o /path/to/output*

**manta (v1.6)**

*configManta.py --normalBam /path/to/normal_sample.bam --tumorBam /path/to/tumor_sample.bam --reference /path/to/ref.fa --runDir /path/to/output*

**nanomonsv (v0.7.2)**

*nanomonsv parse /path/to/tumor_sample.bam /path/to/output/test_tumor*

*nanomonsv parse /path/to/normal_sample.bam /path/to/output/test_normal*

*nanomonsv get /path/to/tumor_sample.bam /path/to/normal_sample.bam /path/to/ref.fa --control_prefix /path/to/output/test_tumor --control_bam /path/to/output/test_normal*

**Severus (v1.5)**

*python severus.py --target-bam /path/to/tumor_sample.bam --control-bam /path/to/normal_sample.bam --out-dir /path/to/output -t 16 --phasing-vcf /path/to/phased_vcf --vntr-bed vntr_bed*

The “*min-support*” parameter of minimum supporting reads note: For the HG002 sample’s HiFi 48×/20×/10×/5× data, “*min-support*” were set to 4/2/1. For the HG002 sample’s ONT 47×/20×/10×/5× data, “*min-support*” were set to 10/4/3/2. For the HG003/HG004 sample’s HiFi and ONT data, “*min-support*” were set to 4 and 10. For the HG005/HG006/HG007 sample’s HiFi and ONT data, “*min-support*” were set to 3 and 10. For rare SVs simulated HiFi and ONT data, “*min-support*” were set to 4 for both. For HG008 and HCC_1395_ sample’s HiFi data, “min-support” were set to 3 and 4.

2	Command lines of SV call eval

*bgzip /path/to/variants.vcf && tabix /path/to/variants.vcf.gz;*

*For germline SV benchmark: truvari bench -f /path/to/ref.fa -b /path/to/ground_truth.vcf.gz -o /path/to/tools --sizemin 50 --sizefilt 50 --passonly -p 0.00 -P 0.5 -r 1000 -c /path/to/variants.vcf.gz --includebed /path/to/include.bed*

*For somatic SV benchmark: truvari bench -f /path/to/ref.fa -b /path/to/ground_truth.vcf.gz -o /path/to/tools --pick multi --sizemin 50 --sizefilt 50 --sizemax -1 --passonly -p 0.00 --dup-to-ins --typeignore -c /path/to/variants.vcf.gz --includebed /path/to/include.bed*

*python /minda/minda.py truthset --base /path/to/ground_truth.vcf.gz --vcfs *.variants.vcf.gz --out_dir minda_multimatch --multimatch --bed /path/to/include.bed*

3	Command lines for alignment

**minimap2 (v2.17-r941)**

*minimap2 /path/to/ref.fa /path/to/sample.fasta -x map-hifi --MD -Y -o /path/to/sample.sam -R ‘@RG\tID:hg2’ -t 64 && samtools view -@ 64 -buS /path/to/sample.sam | samtools sort -@ 64 -O bam -T ./ - > /path/to/sample.bam*

**pbmm2 (v1.17.0)**

*pbmm2 align /path/to/ref.fa /path/to/sample.fasta /path/to/sample.bam --sort --num-threads 64*

**minigraph (v0.21-r606)**

HiFi: *minigraph -t128 -cxasm --vc --secondary yes /path/to/pangenome.gfa /path/to/sample.fasta > /path/to/sample.gaf*

ONT: *minigraph -t128 -cxlr --vc --secondary yes /path/to/pangenome.gfa /path/to/sample.fasta > /path/to/sample.gaf*

For miniSV (stable segment coordinate): *minigraph -t128 -cx asm(lr) /path/to/pangenome.gfa /path/to/sample.fasta > /path/to/sample.gaf*

4	Command lines for down-sample

HiFi-20×: *samtools view -@ 64 -bS -s 0.417 /path/to/sample.bam > /path/to/sample_20*×*.bam*

HiFi-10×: *samtools view -@ 64 -bS -s 0.208 /path/to/sample.bam > /path/to/sample_10*×*.bam*

HiFi-5×: *samtools view -@ 64 -bS -s 0.104 /path/to/sample.bam > /path/to/sample_5×.bam*

ONT-20× and ONT-20×_replicate: *samtools view -@64 -bS -s 0.426 /path/to/sample.bam >*

*/path/to/sample_20×.bam && samtools view -@64 -bS -s 0.426 /path/to/sample.bam –subsample-seed 99 > /path/to/sample_20×_replicate.bam*

ONT-10× and ONT-10×_replicate: *samtools view -@64 -bS -s 0.213 /path/to/sample.bam > /path/to/sample_10×.bam && samtools view -@64 -bS -s 0.213 /path/to/sample.bam –subsample-seed 99 > /path/to/sample_10×_replicate.bam*

ONT-5× and ONT-5×_replicate: *samtools view -@64 -bS -s 0.106 /path/to/sample.bam > /path/to/sample_5×.bam && samtools view -@64 -bS -s 0.106 /path/to/sample.bam –subsample-seed 99 > /path/to/sample_5×_replicate.bam*

5	Command lines for reads simulation of rare SV

**PBSIM3 (v3.0.4)**

ONT: *pbsim --strategy wgs --method qshmm --qshmm R103.model --depth 20 --genome /path/to/sample.fasta*

HiFi: *pbsim --strategy wgs --method sample --sample /path/to/sample.fastq --depth 20 --genome /path/to/sample.fasta*
